# Direct Imaging and Identification of Proteoforms up to 70 kDa from Human Tissue

**DOI:** 10.1101/2021.12.07.471638

**Authors:** Pei Su, John P. McGee, Kenneth R. Durbin, Michael A. R. Hollas, Manxi Yang, Elizabeth K. Neumann, Jamie L. Allen, Bryon S. Drown, Fatma Ayaloglu Butun, Joseph B. Greer, Bryan P. Early, Ryan T. Fellers, Jeffrey M. Spraggins, Julia Laskin, Jeannie M. Camarillo, Jared O. Kafader, Neil L. Kelleher

**Affiliations:** Departments of Molecular Biosciences, Chemistry, Chemical and Biological Engineering, and the Feinberg School of Medicine, Northwestern University, Evanston, IL, USA; Department of Chemistry, Purdue University, West Lafayette, IN, USA; Department of Biochemistry and Mass Spectrometry Research Center, Vanderbilt University, Nashville, TN, USA; Proteomics Center of Excellence, Evanston, IL, USA; Department of Chemistry, Cell and Development Biology, Vanderbilt University, Nashville, TN, USA; Department of Biochemistry and Molecular Genetics, Feinberg School of Medicine, Northwestern University, Chicago, IL, USA

## Abstract

Imaging of proteoforms in human tissues is hindered by low molecular specificity and limited proteome coverage. Here, we introduce proteoform imaging mass spectrometry (PiMS), which increases the size limit for proteoform detection and identification by 4-fold compared to reported methods, and reveals tissue localization of proteoforms at <80 μm spatial resolution. PiMS advances proteoform imaging by combining ambient nanospray desorption electrospray ionization (nano-DESI) with ion detection using individual ion mass spectrometry (I^2^MS). We demonstrate the first proteoform imaging of human kidney, identifying 169 of 400 proteoforms <70 kDa using top-down mass spectrometry and database lookup from the human proteoform atlas, including dozens of key enzymes in primary metabolism. PiMS images reveal distinct spatial localizations of proteoforms to both anatomical structures and cellular neighborhoods in the vasculature, medulla, and cortex regions of the human kidney. The benefits of PiMS are poised to increase proteome coverage for label-free protein imaging of tissues.

**Teaser:** Nano-DESI combined with individual ion mass spectrometry generates images of proteoforms up to 70 kDa.

## Introduction

Proteoforms are the protein-level products of gene expression and post-translational modifications functioning as key effectors in human health and disease (*1, 2*). In addition to understanding of their molecular compositions, interactions, and biological function, comprehensive characterization of the human proteoform landscape also requires mapping of their spatial distributions in human tissues and organs (*3*). Protein-level imaging using antibody-based optical microscopy has revealed distinct cell types, functional tissue units, and subcellular structures (*4*). These techniques employ enzymes, metals, and fluorophores as reporters to obtain high-resolution maps of protein targets in tissues (*5*). In recent years, highly multiplexed antibody-based imaging assays such as CODEX (*6, 7*), IBEX (*8*), and Cell-DIVE (*9*) have drastically increased the number of protein targets that can be probed in a single experiment. Alternatively, mass spectrometry (MS)-based imaging assays, including imaging mass cytometry (IMC) and multiplexed ion beam imaging (MIBI), utilize antibodies labeled with rare earth metals to detect protein localization for up to ~60 protein targets at once (*10, 11*). Despite the significant advances in spatial resolution and sensitivity, antibody-based approaches require prior knowledge of the protein targets and do not provide proteoform-level information (*12, 13*).

MS-based top-down proteomics (*14, 15*) has been widely used for proteoform characterizations (*16*). Modern MS instrumentation has reached the sensitivity for spatially-resolved top-down proteomics suitable for imaging experiments (*17, 18*). Matrix-assisted laser desorption/ionization (MALDI) is widely used for protein imaging (*19, 20*) due to the broad mass range of proteome sampling (*21–23*). However, MALDI predominantly generates singly-charged ions, which gives limited fragment information for direct top-down identification of intact proteins (*24*). This challenge may be addressed using matrix-assisted laser desorption electrospray ionization (MALDESI), which combines MALDI with extractive ESI to generate multiply-charged ions of peptides and proteins extracted from tissues (*25*). Alternatively, multiply-charged protein ions may be generated using liquid extraction-based ambient ionization methods (*26*) including desorption electrospray ionization (DESI) (*27*), liquid extraction surface analysis (LESA) (*28*), and nanospray desorption electrospray ionization (nano-DESI) (*29*). These techniques are particularly advantageous in top-down analysis of intact proteoforms in the imaging mode. Among these techniques, nano-DESI that utilizes a sub-nanoliter dynamic liquid bridge as a sampling probe enables imaging of biomolecules in tissues with a spatial resolution down to 10 μm (*30, 31*).

One major challenge in proteoform imaging using liquid extraction-based techniques is the detection of low-abundance, high mass proteoforms in the congested MS spectra produced by ionizing complex mixtures of biomolecules extracted from the sample. Until now, imaging and identification of intact proteoforms directly from tissue has been limited to <20 kDa species (*27, 28, 32, 33*), with one report leading to the identification of subunits from a 43 kDa trimeric protein complex (*34*). Here, we address this challenge using individual ion mass spectrometry (I^2^MS). I^2^MS is a new Orbitrap-based charge detection technique (*35–37*) utilizing individual ions. Using this workflow yields a 10-20-fold enhancement in resolving power and 500× dilute sample compatibility compared to traditional ensemble MS techniques (*38*). In particular, we combine nano-DESI ionization (*33*) with I^2^MS (*39*) to create proteoform imaging mass spectrometry (PiMS) for tissue imaging and direct identification of proteoforms up to ~70 kDa. We show ~400 isotopically-resolved proteoform assignments from human kidney and confidently identify proteoforms up to 53 kDa using MS/MS, illuminating differences in kidney architecture from the medulla, cortex, and vasculature. Incorporating I^2^MS, PiMS yielded 169 proteoform assignments/identifications at 80 μm spatial resolution and demonstrates the potential to visualize the proteinaceous structures comprising human tissues.

## Results

### Overview of PiMS workflow

Proteoform imaging mass spectrometry (PiMS) illustrated in Fig. 1 combines nanospray desorption electrospray ionization (nano-DESI) imaging (*33*) with data acquisition and processing for individual ions (*39*). Specifically, we perform nano-DESI line scans on tissue, during which proteoforms are sampled as multiply-charged ions distributed across multiple charge states (Fig. 1, top left). Instead of unresolved protein signals typically observed for ensembles of ions, we detect charge-assigned individual protein ions as a function of location on the tissue section thereby generating mass spectral features of the individual proteoforms with resolution of their ^13^C isotopic peaks in each pixel of the imaging data (fig. S1). This allows for confident assignment of proteoform masses with better than 2 parts-per-million accuracy at one sigma in each pixel of the imaging data (Fig. 1, middle left). Beyond proteoform-specific images (Fig. 1, bottom left), molecular identification is achieved using either direct top-down MS/MS off the tissue (Fig. 1, top right) or database searching from known proteoforms of the intact masses (Fig. 1, bottom right).

**Fig. 1.**
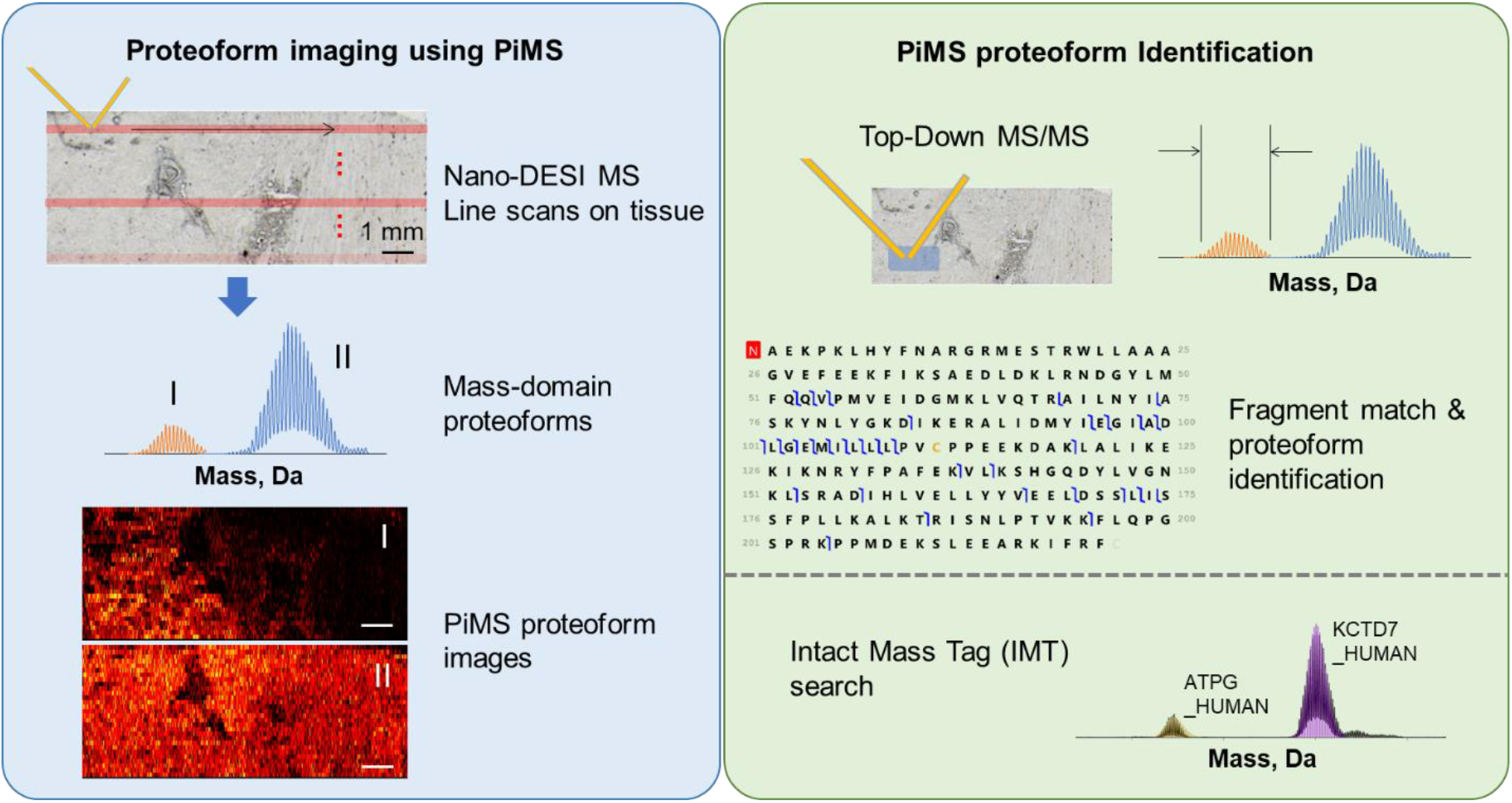
Illustration of the PiMS workflow for proteoform imaging and identification. The left panel shows the scanning approach (top), detection of proteoforms in the mass domain (middle) and image reconstruction (bottom). The right panel depicts the two approaches to identify proteoforms using either direct fragmentation of proteoform ions and spectral readout by individual ion MS/MS (top) or database lookup of accurate mass values (IMT, bottom).

### Human Kidney Proteoforms Detected by PiMS

We used PiMS to examine the proteoforms and their localizations in a 10 μm-thick human kidney tissue section. Encouragingly, we immediately expanded the mass detection range for proteoform imaging to >70 kDa. Fig. 2A shows the full PiMS spectrum from 5-72 kDa from a sum of 16500 MS scans (~8 million single ions). This spectrum contains ~400 proteoform masses above 0.1% relative abundance that are isotopically resolved. A complex group of proteoforms in the 68-80 kDa range were observed, but not individually resolved (fig. S2). Spectral attributes include a dynamic range of ~200 (using S/N 3 as the limit of detection) and a mass resolution (*m/Δm*) of ~100,000. PiMS images can be constructed for any of the 242 proteoform masses with relative abundance above 1% in the full PiMS spectrum of Fig. 2A.

**Fig. 2.**
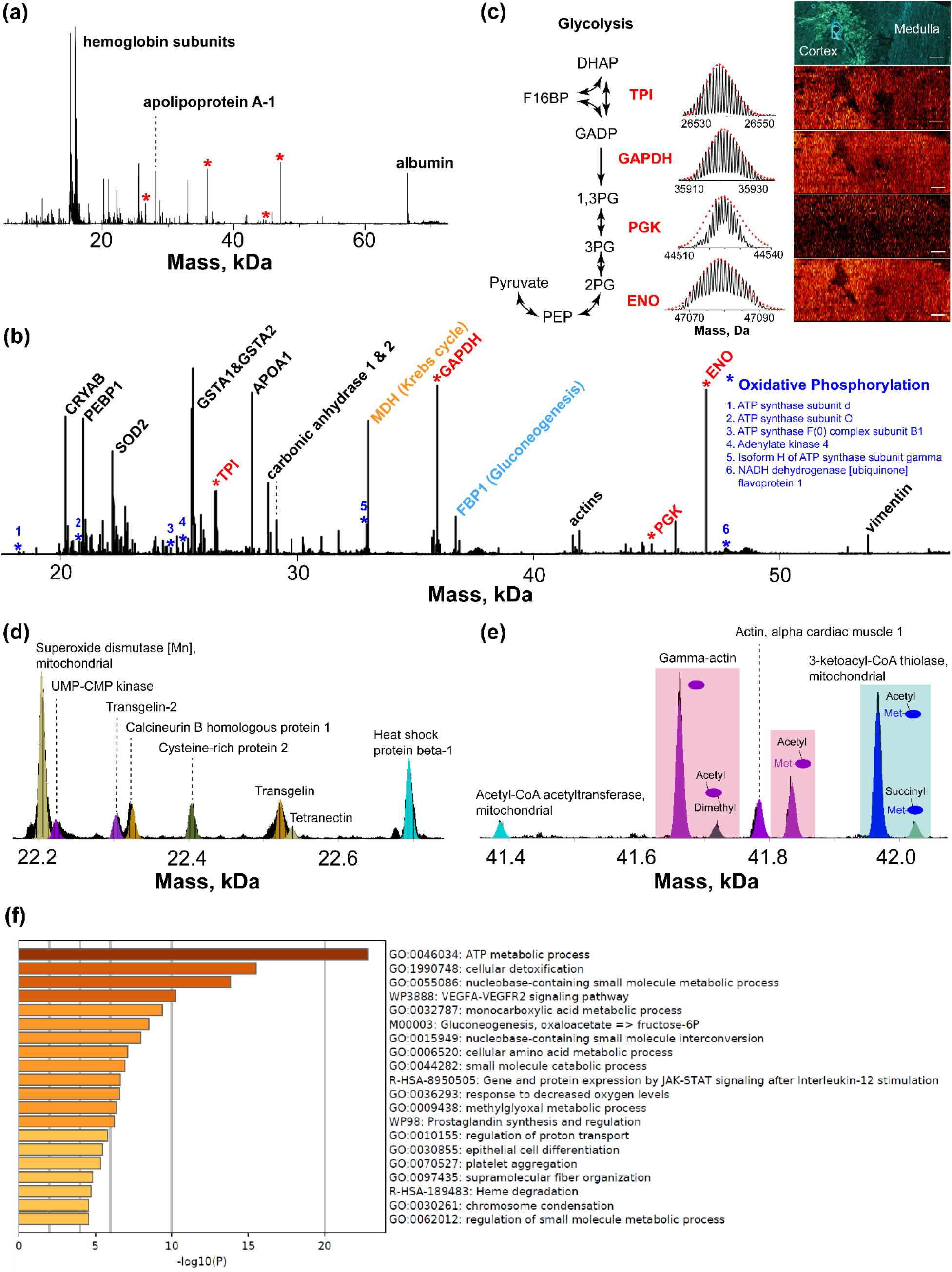
Sum of mass domain spectrum obtained from PiMS of human kidney. (a) Full scale PiMS spectrum in 5-72 kDa summed from 16500 MS scans; regions occupied by abundant blood proteins (hemoglobin subunits, albumin) are labeled in the spectrum; red asterisks denote the key glycolytic enzymes found in the spectrum. (b) PiMS full spectrum in the 18-56 kDa range zoomed in from (a); major proteins identified using a variety of approaches are labeled in the spectrum; aside from glycolytic enzymes (red asterisks), enzymes involved in a few other major metabolic pathways (Krebs cycle, gluconeogenesis, oxidative phosphorylation) found in PiMS are labeled in the spectrum. (c) Theoretical (red triangles) and experimentally-observed (black trace) isotopic distributions of the four glycolytic enzymes (labeled with red asterisks in (a) and (b)) together with their PiMS images depicted in a schematic diagram of the glycolysis metabolic pathway. The cortex and the medulla of the kidney section imaged are labeled in the autofluorescence image at top. Scale bar: 1 mm. Selected mass range of 22.2-22.7 kDa (d) and 41.4-42.1 kDa (e) showing identified proteoforms (spectrum in black, theoretical isotopic distributions in color). (f) GO analysis of biological pathways found enriched in the 169 identified proteoforms from the PiMS experiment shown in the -log(10)P scale.

To identify these proteoforms, we manually annotated the full PiMS spectrum shown in Fig. 2A based on the results of an intact mass tag (IMT) search against a custom database. In this approach, we compared the shape and mass accuracy of the isotopic distributions of the proteoforms in the PiMS spectrum with the theoretical proteoforms in the database. The database was constructed from the top 500 most abundant proteins identified in a bottom-up proteomics study of human kidney tissues (*40*). To maximize the number of proteoform identification, we combined additional matches from top-down identification (details discussed in the next section) and manually inspected post-translational modifications (PTMs) recorded in the Swiss-Prot database (*40*). As a result, we manually annotated 169 proteoforms in the entire mass range using a ±5 ppm mass tolerance (list of proteoforms shown in table S1). Fig. 2D and 2E show two zoomed regions of the full PiMS spectrum with theoretical proteoform matches highlighted in color. Fig. 2D shows various unmodified proteoforms in the mass range of 22.2-22.7 kDa captured by the custom database, demonstrating that the search included a variety of proteins in the kidney proteome. The mass range of 41.4-42.1 kDa shown in Fig. 2E contains proteoforms of gamma-actin (highlighted in pink) and 3-ketoacyl-CoA thiolase (highlighted in blue) exhibiting diverse PTMs and their combinations. Aside from the monoacetylated and dimethylated proteoform of gamma-actin identified by top-down MS, other modified proteoforms in the displayed mass range were manually annotated. Clearly, further investigation is needed to estimate the false discovery rate for automated identification in PiMS, including the use of Bayesian priors.

We annotated the most abundant proteins in Fig. 2A and Fig. 2B to demonstrate the portion of the human kidney proteome captured by PiMS. Not surprisingly, blood proteins (hemoglobin subunits, apolipoprotein A-1, and albumin, Fig. 2A) were found at highest abundance due to the highly vascularized nature of the kidney. Meanwhile, we captured many proteins prevalent in cellular pathways that are naturally abundant in human cells. In Fig. 2B, we labeled the most abundant proteins to give a brief overview of the molecular functions and biological pathways observed in PiMS. From low to high mass range shown in Fig. 2B, we found molecular chaperone (alpha-crystallin B chain, CRYAB), signaling modulator (phosphatidylethanolamine-binding protein 1, PEBP1), proteins for cellular detoxification (superoxide dismutase [Mn], mitochondrial, SOD2, glutathione S-transferase A1&A2, GSTA1&GSTA2) and homeostasis (carbonic anhydrase 1&2), and structural proteins (actins, vimentin). Moreover, proteins participating in central metabolic pathways are dominant. In particular, we found key enzymes in Krebs cycle (malate dehydrogenase, MDH) and gluconeogenesis (fructose-1,6-bisphosphatase 1, FBP1), and 27 subunits of protein complexes in the electron transport chain of oxidative phosphorylation (blue asterisks for 6 subunits in 18-56 kDa mass range, others recorded in table S1). More intriguingly, four key enzymes in glycolysis (triosephosphate isomerase, TPI, glyceraldehyde-3-phosphate dehydrogenase, GAPDH, phosphoglycerate kinase, PGK, and alpha-enolase, ENO) were imaged and identified (Fig. 2C, red asterisks in Fig. 2A and 2B). PiMS images of the four detected glycolytic enzymes were shown in Fig. 2C with largely even distributions across the entire tissue section spanning from the kidney medulla to the cortex, confirming the presence of glycolysis in many of the kidney cell types. A more complete investigation of the biological pathways found for the 169 identified proteoforms is demonstrated by a Gene Ontology (GO) analysis shown in Fig. 2F, which indicates that cellular metabolic processes is the predominant biological pathway observed in the PiMS experiment.

### Top-Down Characterizations of Kidney Proteoforms in PiMS

Human kidney proteoforms were characterized using tandem MS (MS/MS). Direct fragmentation of protein ions >20 kDa is challenging due to the low abundance of their fragment ions, particularly larger fragments (>15 kDa) that result from cleavage in the middle of the protein sequence (*41*). To overcome this limitation, we employed I^2^MS for the readout of top-down fragmentation spectra to capture those large fragment ions typically buried under the noise level in ensemble MS/MS experiments (*41*). In particular, we selected 20-50 kDa proteoforms observed at >4% relative abundance for on-tissue MS/MS aiming for where they are in highest abundance on the tissue section. For each target proteoform, we selected a <0.8 *m/z* wide isolation window corresponding to the most abundant charge state of the proteoform obtained from PiMS data at MS^1^-level (Fig. 1, right and Materials and Methods section). By matching the originally observed intact mass with the subsequent fragmentation data, we confidently identified 21 proteoforms >20 kDa with E-values ranging from 10^-12^ to 10^-160^ (Table 1 and fig. S7a-u).

**Table 1.**
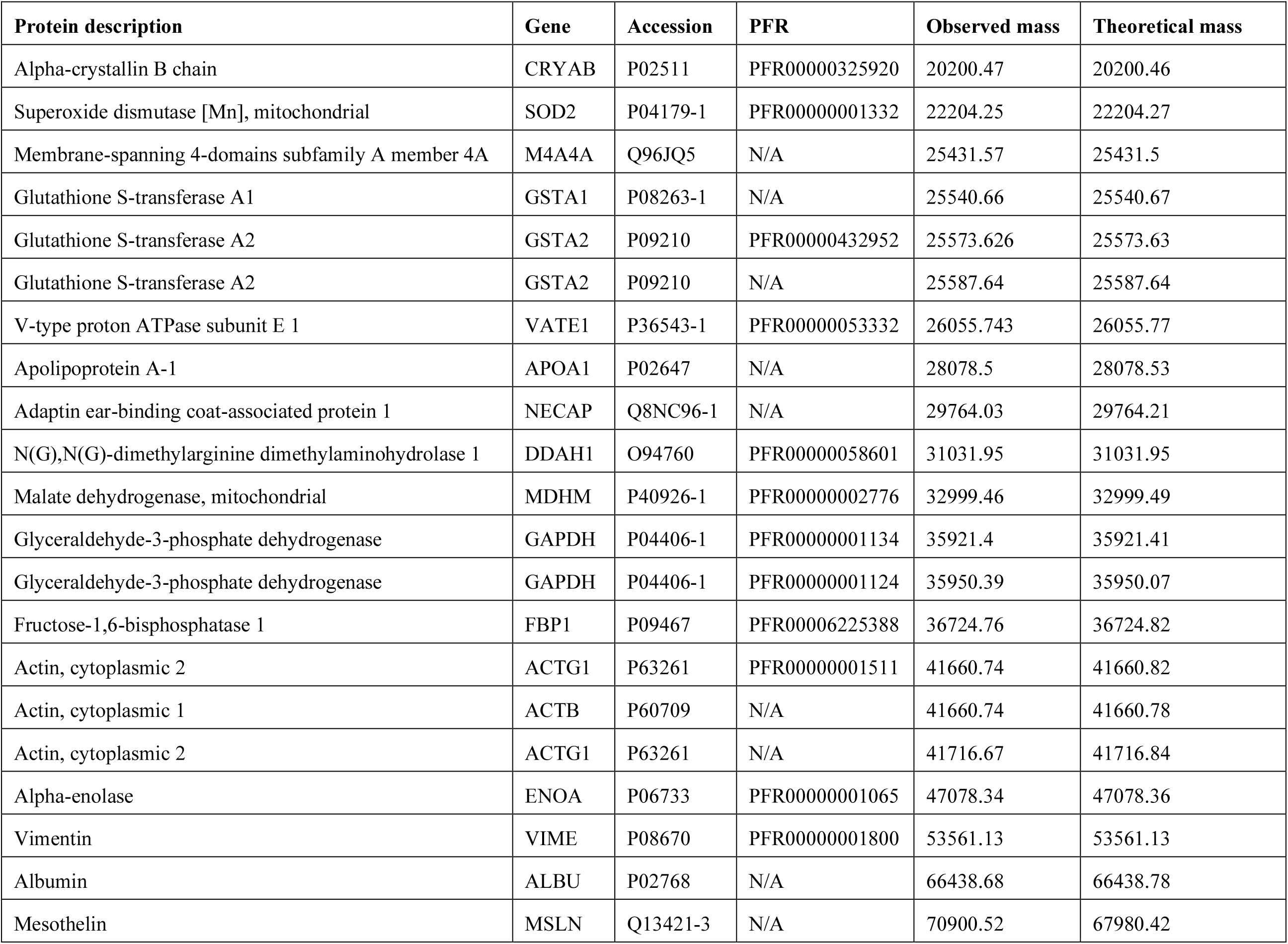
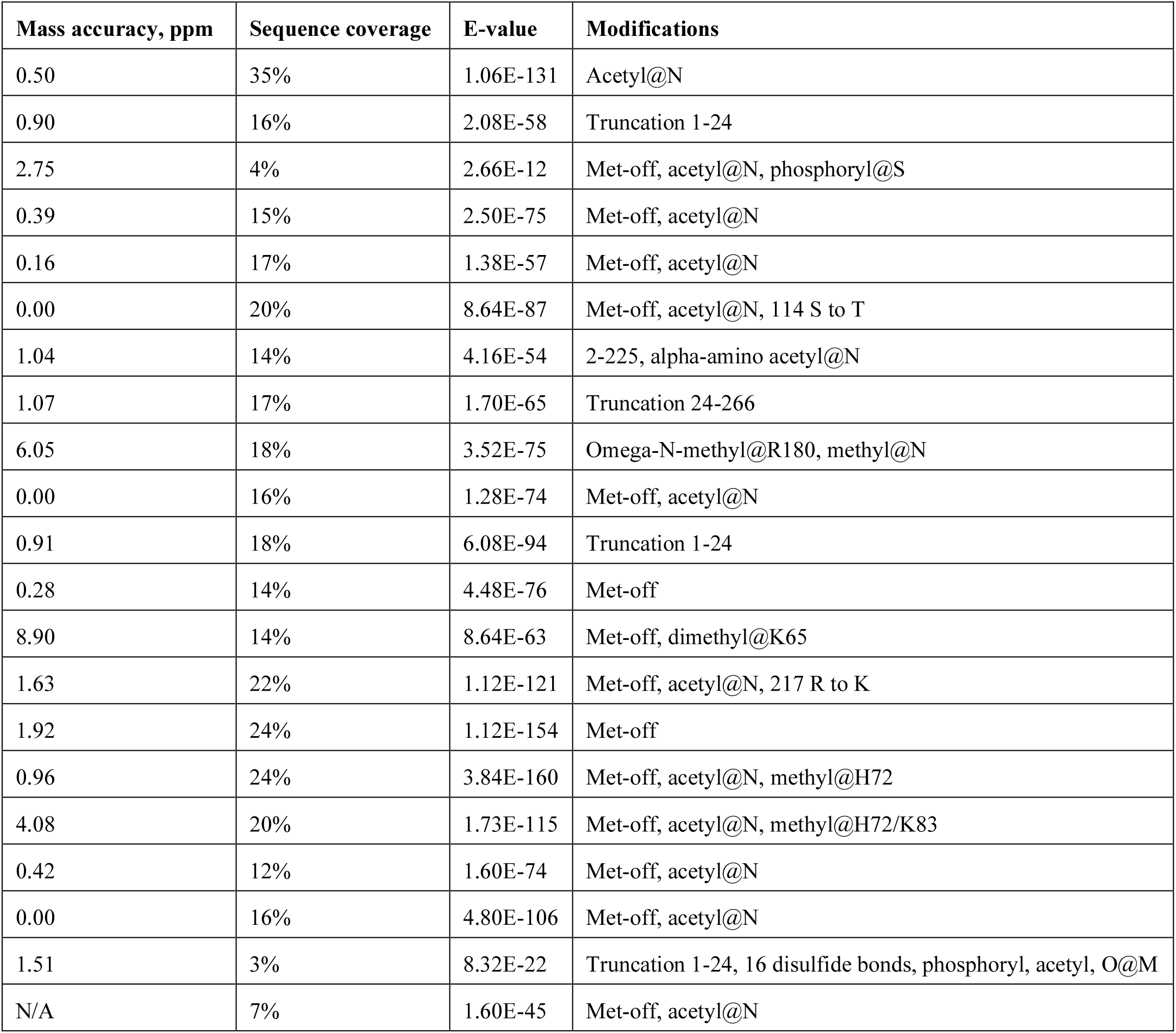
Proteoforms identified by on-tissue MS/MS.

In Fig. 3, we highlight two representative >20 kDa proteoforms confidently identified by MS/MS. The 11-14.5 kDa region of the fragmentation spectrum of monoacetylated glutathione S-transferase A1 (GSTA1, 25,542 Da, Fig. 3A) contains abundant complementary fragments of a 14-amino-acid-long sequence tag (B95-B108, Y113-Y126), contributing to the confident identification of this proteoform (Graphical Fragment Map, GFM, shown in Fig. 3A). PiMS image of GSTA1 proteoform in Fig. 3A shows that this proteoform is localized to the kidney cortex region. On the higher mass end, vimentin (53,530 Da) was also identified (fragment spectrum and GFM shown in Fig. 3B), with 63 isotopically-resolved >15 kDa fragment ions above 1% relative abundance matching sequence fragments of vimentin (a few representative ones shown in Fig. 3B). PiMS image of vimentin (Fig. 3B) showing its localization to the vasculature also confirms the identity of the proteoform (*42*).

**Fig. 3.**
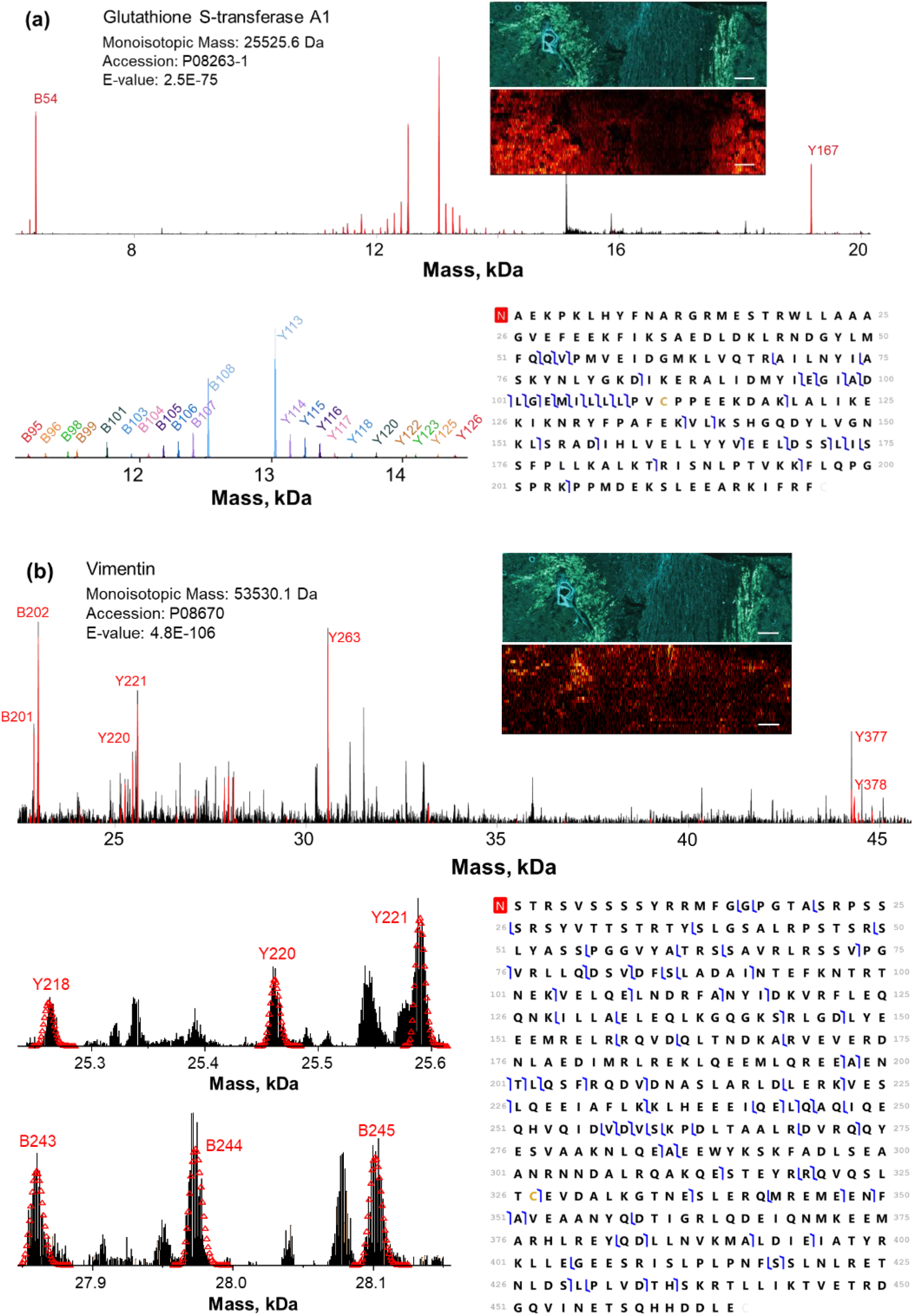
On-tissue identification of proteoforms by MS/MS. Two representative human kidney proteoforms, glutathione S-transferase A1 (a) and vimentin (b). The PiMS images of the two proteoforms are shown on the top right of each panel along with an autofluorescence image of an adjacent section as a reference. On the bottom left of each panel, expanded regions of the fragment spectrum are displayed with the major matching fragment ions annotated. GFMs are shown on the bottom right of each panel. Scale bar: 1 mm.

The identification of the >60 kDa proteoforms by MS/MS is increasingly challenging. We obtained 3% sequence coverage for a ~66.4 kDa proteoform putatively identified as albumin (fig. S7t). Poor sequence coverage obtained for this proteoform is likely due to presence of 17 disulfide linkages known to occur in albumin, thereby making PTM localization challenging. In one attempt to identify a proteoform centered at 70,900 Da, we found that the precursor proteoform with the best database retrieval score was mesothelin isoform 2 (fig. S7u). The deviation in precursor mass (67,938 Da compared to 70,900 Da) may be attributed to modifications and/or isoform expression, which are not captured in the database. Despite these challenges, we were able to readily identify 21 human proteoforms ranging from 20 to 70 kDa in molecular mass using top-down MS in PiMS.

### Creation of a Kidney Proteoform Map

PiMS images allow for direct visualization of sub-mm anatomical structures and functional tissue units of human kidney sections with proteoform-level precision. PiMS of kidney tissue containing cortex, medulla, and vasculature regions shows distinct differences in the distribution of proteoforms across these vastly different anatomical regions (Fig. 4, optical image shown in Fig. 4A). The identification of kidney internal structures was supported by autofluorescence microscopy (Fig. 4B) (*43*) and periodic acid-Schiff (PAS) staining histology (Fig. 4C).

**Fig. 4.**
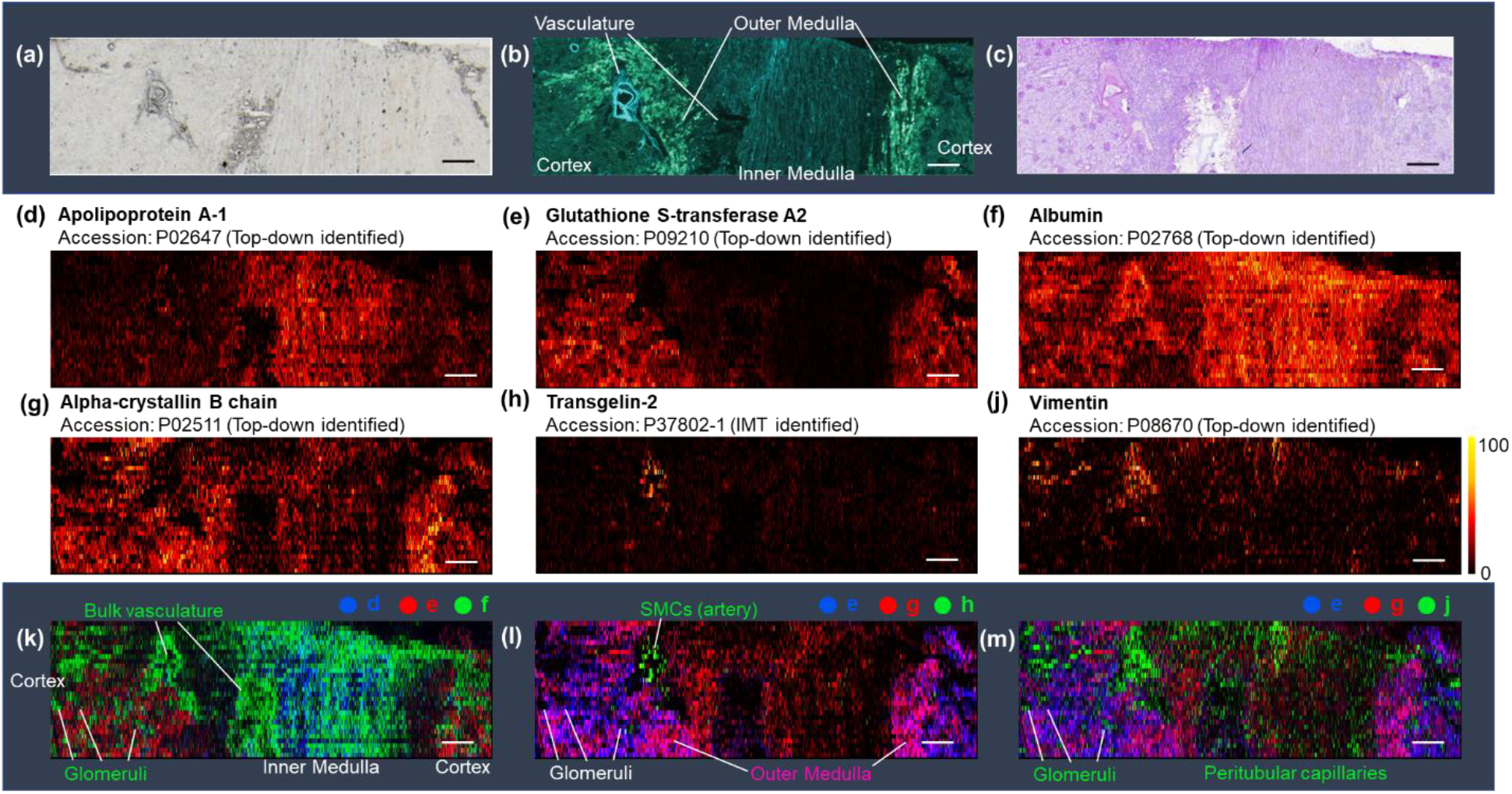
Kidney Proteoform Maps. Optical (a) and autofluorescence (b) images of adjacent human kidney sections containing the cortex, medulla, and vasculature regions; (c) PAS staining of an adjacent section from the same kidney; (d)-(j) PiMS images of individual proteoforms that selectively illuminate different anatomical regions and cellular neighborhoods, the name and UniProt accession of the proteoforms are depicted next to the images with their color scale. Composite image of (k) apolipoprotein A-1 (blue, d), glutathione S-transferase A2 (red, e), and albumin (green, f); (l) glutathione S-transferase A2 (blue, e), alpha-crystallin B chain (red, g), and Transgelin-2 (green, h); and (m) glutathione S-transferase A2 (blue, e), alpha-crystallin B chain (red, g), and vimentin (green, j). Scale bar: 1 mm.

PiMS images of proteoforms in the kidney show distinct localizations (Fig. 4D-J). Apolipoprotein A-1 (Fig. 4D) and GSTA2 (Fig. 4E) were found to be enhanced in the medulla and cortex regions, respectively. Blood-abundant albumin (Fig. 4F) and alpha-crystallin B chain (Fig. 4G), a molecular chaperone, were ubiquitously expressed in most of the regions of the kidney. Additionally, transgelin-2 (Fig. 4H) and vimentin (Fig. 4J) were abundant only in highly-focused regions near the artery.

### Assessing Proteoform Biology: Differences in Space and Molecular Composition

Next, we created composite PiMS proteoform images to enable more efficient readout of anatomical regions and functional tissue units in the kidney. Using the abovementioned single PiMS images, we created a series of tricolor composite PiMS images (Fig. 4K-M). By combining images of apolipoprotein A-1 (Fig. 4D, blue), GSTA2 (Fig. 4E, red) and albumin (Fig. 4F, green), we highlighted vascular regions within the kidney (Fig. 4K, labeled as bulk vasculature). This confirms the highly vascularized inner medulla, along with distinct points of vascularization in the cortex aligning with glomeruli. Fig. 4L is a composite PiMS image of transgelin-2 (Fig. 4H), GSTA2 (Fig. 4E, blue) and alpha-crystallin B chain (Fig. 4G, red). Using this image, we highlight the large artery via the specific localization of transgelin-2 (Fig. 4L, green). Previous studies have shown the specific localization of transgelin-2 to the smooth muscle cells (SMCs). This observation is consistent with the abundance of SMC in arteries whereas veins are mainly comprised of stromal cells (*44*). Finally, the combination of the spatial distribution of vimentin (Fig. 4J, green), GSTA2 (Fig. 4E, blue), and alpha-crystallin B chain (Fig. 4G, red) provides the best outline of the vasculature in the tissue section. Vimentin is a filamentous protein found in most of the blood vessels and connective tissues. The localization of vimentin to the bulk blood vessel regions observed by PiMS is consistent with this. A majority of the scattered vimentin spots observed in the cortex fit well into the dark spots corresponding to glomeruli. Moreover, vimentin is found in many scattered locations in the inner medulla region. These illuminated spots correspond to the spatially-dispersed peritubular capillaries, which form a complex three-dimensional network in the medulla and become dispersed when the tissue is cross-sectioned.

Moreover, a major advantage of PiMS over antibody-based imaging approaches lies in the ability to determine the molecular composition of proteoforms in an untargeted fashion. Antibody-based imaging approaches do not distinguish different proteoforms of a single protein, whereas PiMS can capture sequence differences and modifications. Subtle differences of protein sequences and modifications become especially challenging to detect for high-mass proteins. Direct top-down identification of >20 kDa proteoforms from tissue enabled by PiMS allows for the characterization of highly similar kidney protein isoforms that originate from allelic coding single nucleotide polymorphisms (cSNPs).

Three major proteoforms of N-terminal acetylated GST subunits, GSTA1 and GSTA2, were observed in kidney tissue with localizations to the cortex region (Fig. 5B, 5D, and 5E). Fig. 5C shows the mass domain spectrum of GSTA1 and GSTA2 proteoforms. A single proteoform was detected from GSTA1 (25,542 Da), showing two alleles with the same sequence. In contrast, we identified two GSTA2 proteoforms, the canonical form at 25,573 Da, and another form at 25,587 Da representing a 14 Da mass shift. Both proteoforms were observed at similar abundances with highly similar tissue localization, which is characteristic of non-specific biallelic tissue expression resulting from a cSNP (Fig. 5D and 5E). Fragmentation data was able to localize the 14 Da mass shift to the region between Pro110 and Gln113 from the N-terminus (regions highlighted in light red, Fig. 5D and 5E and fig. S7d). A UniProt search shows a Ser111 → Thr natural variant of GSTA2 (highlighted in red), confirming that the 14 Da mass shift corresponds to a proteoform resulting from a common cSNP (allele frequency >40% according to dbSNP entry No. rs2180314). This exemplifies the power of PiMS to the probing of gene expression in tissues directly at the proteoform level, which is complementary to genomic and transcriptomic predictions.

**Fig. 5.**
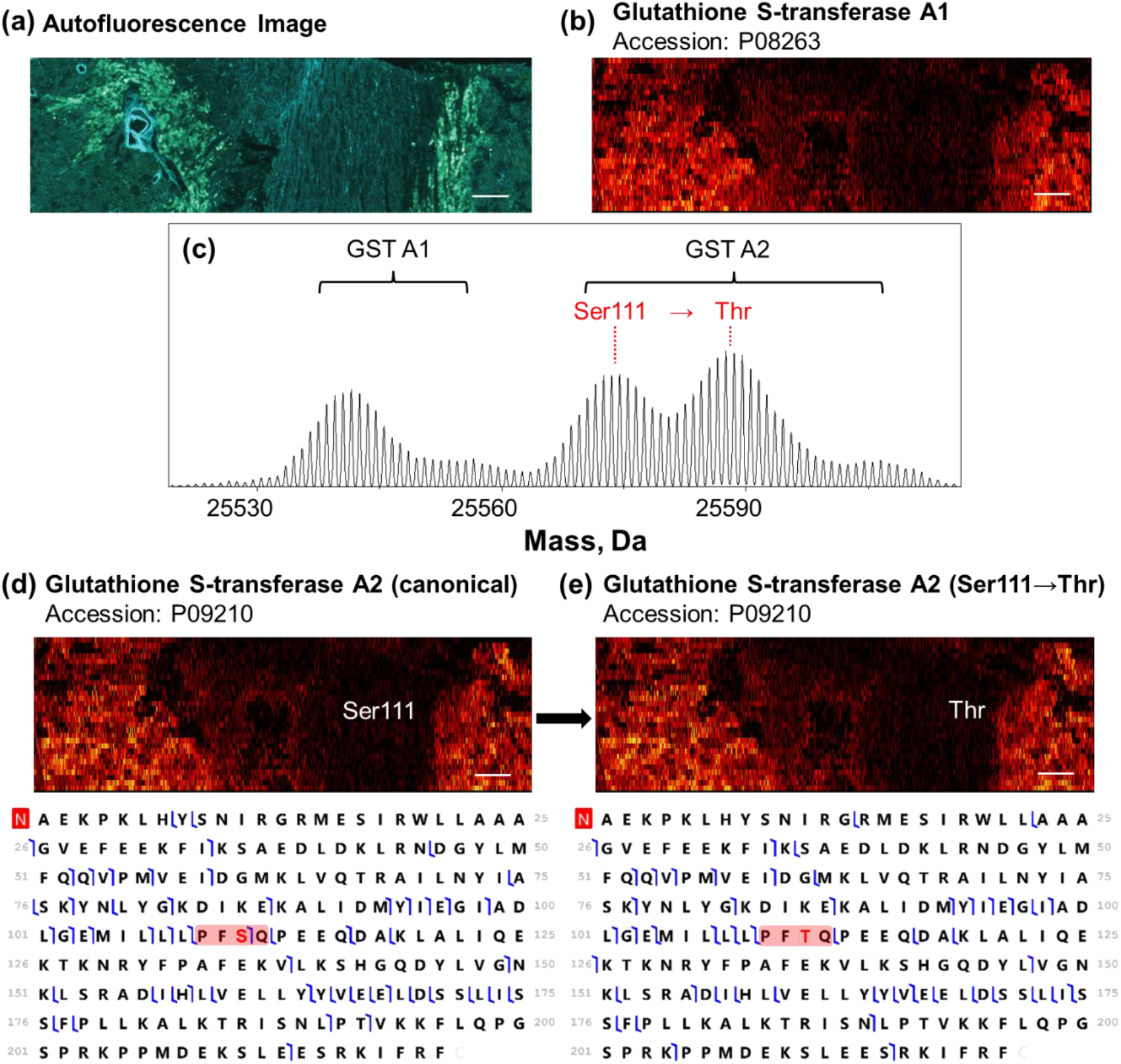
Mass spectrometric detection of gene differences. (a) Autofluorescence and PiMS images of alpha GST enzymes of the human kidney section (b, d, and e). Mass domain PiMS spectra of ~25.6 kDa range (c) shows GSTA1 & GSTA2 enzyme proteoforms. The GFMs of the two proteoforms of GSTA2 from a known biallelic cSNP are shown in (d) and (e) below the PiMS images. The sequence variation of the two cSNP GSTA2 proteoforms are highlighted in the GFMs in red.

## Discussion

Imaging methods have boomed in recent years. Highly multiplexed affinity reagent-based methods allow for the detection of more than 50 protein targets in a single assay (*6*). The increased throughput of these approaches has provided an opportunity to develop comprehensive maps of proteins in human tissues at a rapid pace (*4, 8, 9*). However, antibody-based imaging approaches do not distinguish different proteoforms of a single protein. The development described in this study addresses this challenge and enables imaging protein isoforms from different gene family members or allelic cSNPs in biological tissues.

PiMS presents low-bias to the protein masses compared to multiplexed imaging assays, with the ability of capturing a majority of the abundant cytosolic proteins in cells. In fact, while performing IMT search of PiMS data against a database containing 100 most abundant proteins from bottom-up proteomics of human kidney tissue, 70 protein candidates were detected by PiMS. This provides an opportunity of using PiMS to delve into the major biological pathways in tissues in a spatially-resolved manner. In the GO analysis result shown in Fig. 2F, a variety of central cellular metabolic pathways were found enriched by PiMS. Notably, we were able to detect and image four key enzymes in glycolysis and 27 subunits of the complexes in the respiratory electron transport chain, which are critical component of ATP metabolic process, the highest enriched pathway.

Another advantage of whole proteoform measurement is the robust detection of diverse types of post-translational modifications with low bias and knowledge of their stoichiometry. For example, two proteoforms of GAPDH were identified in the kidney dataset, both missing the start methionine and one containing a dimethylation site at Lys65. While both of these post-translational modifications are known (*45, 46*), the dimethylation of Lys65 was shown to be very low in abundance and inconsistently observed (*46*). Our data indicates that the dimethylated proteoform is about ~5% of the total GAPDH in human kidney. The function of the dimethylation is unknown, but its colocalization with the unmethylated proteoform suggests a possible role of the modified GAPDH proteoform in dimerization, which may contribute to its catalytic properties in glycolysis (*47*).

In conclusion, we present the PiMS approach that combines nano-DESI with I^2^MS technology, which enables the direct imaging and molecular identification of human kidney proteoforms up to ~70 kDa. This approach increases observable proteoform masses by nearly 4-fold and resolving power by 10-fold compared to prior work (*48*). The development of PiMS opens up exciting opportunities to infuse proteoform knowledge into the multi-omic approaches being evaluated for inclusion into the Human Reference Atlas (*49*). By providing spatial localization of proteoforms to anatomical regions, cell types, and functional tissue units, PiMS promises applications in molecular tissue mapping, biomarker discovery, and disease diagnostics.

## Materials and Methods

### Tissue Preparation

Human kidney tissue sections were prepared according to published protocols (*50*). Mouse brain tissue sections were sectioned at −21°C to a 12 μm thickness using a CM1850 Cryostat (Leica Microsystems, Wetzlar, Germany). Tissue sections were thaw mounted onto glass microscope slides (IMEB, Inc Tek-Select Gold Series Microscope Slides, Clear Glass, Positive Charged) and stored at −80 °C before mass spectrometry imaging analysis.

Human kidney tissue sections were thawed under slight vacuum at room temperature, fixed and desalted via successive immersion in 70%/30%, 90%/10%, and 100%/0% ethanol/H2O solutions for 20 s each, delipidated by 99.8% chloroform for 60s, and dried under slight vacuum right before nano-DESI imaging experiments. These sample preparation steps allow for the *in situ* precipitation of proteins and removal of lipids avoiding suppression of protein signals upon ionization into the mass spectrometer (*19, 51*).

### Nano-DESI Ion Source

A custom-designed nano-DESI source was used for all data acquisition. The experimental details of nano-DESI MSI have been described elsewhere (*29, 30*). Briefly, the nano-DESI probe is comprised of a primary (OD 150 μm, ID 20 μm) and a nanospray capillary (OD 150 μm, ID 40 μm) with the spray side of the nanospray capillary positioned close to the MS inlet. The probe was fabricated using fused silica capillary tubing (Molex, Thief River Falls, MN). A liquid bridge formed at the location where the two capillaries meet is brought into contact with the tissue section for analyte extraction. The liquid bridge is dynamically maintained by solvent propulsion from the primary capillary and instantaneous vacuum aspiration through the nanospray capillary. The extracted analytes are continuously transferred to a mass spectrometer inlet and ionized by ESI. Imaging experiments are performed by moving the sample under the nano-DESI probe in lines. The optimal scan rate is discussed in the next section. The strip step between the line scans was set to 150 μm to avoid overlap between the adjacent line scans. To ensure the stability of the nano-DESI probe during the imaging experiment, we applied a surface tilt angle correction to the tissue sample by defining a three-point plane prior to the imaging experiment (*52*). All samples were electrosprayed under denaturing conditions in a 60%/39.4% acetonitrile/water and 0.6% acetic acid solution compatible with both protein extraction and ionization. All the experiments were performed in positive ionization mode.

### PiMS Conditions and Data Acquisition

PiMS data acquisition was performed in the individual ion mass spectrometry (I^2^MS) mode which has been described previously on a Q-Exactive Plus Orbitrap mass spectrometer (Thermo Fisher Scientific, Bremen, Germany, fig. S1) (*48*). The source conditions on the mass spectrometer were set as follows: ESI voltage: 3 kV; in-source CID: 15 eV; S-lens RF level: 70%; capillary temperature: 325 °C. In particular, rather than collecting typical ensemble ion MS spectra, ion signals were attenuated down to the individual ion regime by limiting the ion collection time in the C-trap before injection into the Orbitrap analyzer. The MS acquisition rate was set at 1 scan every 2 s. During PiMS data acquisition, proteoforms were sampled by a nano-DESI probe producing multiply-charged ions distributed across multiple charge states. To enable downstream I^2^MS analysis, the majority of ions in one detection period was collected in the individual ion regime, corresponding to a singular ion signal at a defined *m/z* (or frequency) value. In particular, rastering line scans on tissue were performed at a reduced rate of 2.5-4 μm/s, which corresponds to a 5-8 μm sampling distance between adjacent pixels (fig. S3). The nano-DESI solvent flow rate was kept at 600 nL/s to allow for efficient extraction and dilution of the proteins. The injection time for a specific set of tissue sections is typically optimized by acquiring line scans on an adjacent section prior to an imaging experiment, and it may vary from 100 to 500 ms (fig. S4-fig. S6). For 10 μm thin sections of human kidney tissue presented in this study, a 300 ms injection time was employed for all the sections from the same subject. We note that for extremely dominant proteoforms (e.g., hemoglobin subunits in this study), “multiple ion events” were commonly observed.

Additional MS instrument conditions in the PiMS experiment are mentioned here: the Orbitrap central electrode voltage was adjusted to −1 kV to improve the ion survival rate under denatured conditions. HCD pressure level was kept at 0.2 (UHV pressure < 2 × 10^-11^ Torr) to reduce collision-induced ion decay within the Orbitrap analyzer without substantial losses in trapping efficiency. Additional relevant data acquisition parameters were adjusted as follows: mass range: 400-2500 *m/z*; AGC mode was disabled and the maximum injection time was held constant at 300 ms; enhanced Fourier transform: off; averaging: 0; microscans: 1. Time-domain data files were acquired at detected ion frequencies and recorded as Selective Temporal Overview of Resonant Ions (STORI) files (*53*).

### PiMS Data Analysis & Image Generation

Ion images were generated using a MATLAB script developed in-house. Mass-domain spectra were constructed by co-adding all individual ions obtained from the entire tissue section. In specific cases where computation power was limited, charge assignment and image construction were performed in sections with an upper limit of 50 million ions per portion. In the first step, all ion signals were subjected to STORI analysis to filter out decayed and multiple ion events. The neutral masses of the protein ions were calculated by:

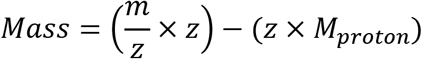

Charge state (*z*) is obtained from the slope of induced image current determined by the STORI analysis (*53*). Accurate charge assignment of each ion was statistically evaluated by comparing the slopes of its isotopologues across different charge states from the entire tissue section. In particular, an iterative voting methodology was employed for filtering out ions with a lower probability score in the process, which allows for the construction of mass-domain isotopic distribution of a proteoform with statistical confidence. In this step, we utilized a Kernel Density Estimation (KDE) approach to convert centroid masses of individual ions to uniform distributions. Accurate masses of the isotopes were obtained from the center of the summed individual ion profiles.

For image generation, ions composing the mass-domain isotopic envelope of a protein were registered back to their spatial origins on the tissue section for PiMS image generation. A ±10 ppm isotopic mass tolerance was used to select individual ions for image generation. A raw image was first generated using absolute ion counts at different x-y locations. In the kidney PiMS images presented in this study, each pixel was constructed from three adjacent MS scans corresponding to ~10 μm × 150 μm area. The raw image was normalized using a total ion count matrix, which accounts for the fluctuation of sampling conditions at different locations.

### Intact Mass Tag (IMT) Search & Gene Ontology (GO) analysis

The summed mass-domain PiMS spectrum was converted to .*mzML* format and processed using a custom version of TD Validator (Proteinaceous, Evanston, IL) implemented with an MS^1^ IMT search function. The PiMS spectrum was shifted by +4 ppm according to the accurate masses of six MS/MS identified proteoforms in the 20-50 kDa mass range. A human protein database constructed from top 500 most abundant proteins in a bottom-up proteomics study of human kidney tissues was used for the search (Table S2). Methionine on/off and monoacetylation were considered as possible proteoform modifications in the database. IMT search was performed with a ±5 ppm mass tolerance. Additional proteoform matches were curated by spectrum inspection and manual annotation of putative modifications recorded in Swiss-Prot human proteome database. The final 169 proteoform matches include MS/MS-identified proteoforms and all IMT-identified proteoforms discussed above.

GO analysis was performed using Metascape (https://metascape.org/) (*54*). Specifically, a list of Entrez Gene ID was retrieved for the 169 identified proteoforms on Uniprot and submitted to Metascape for GO analysis. The result contains the top-level GO biological processes.

### Top-Down Proteomics Data Acquisition & Analysis

Targeted MS/MS experiments were performed on a tissue section adjacent to the imaged section using higher-energy collisional dissociation (HCD). In the first step, a target proteoform was selected in the mass-domain spectrum. We utilized the PiMS image of the target proteoform to select a target area on the section where the proteoform abundance is enhanced. For the selected area, the mass-domain PiMS spectrum was convoluted back to *m/z* domain, from which a proper isolation window that contains predominantly the target proteoform was selected. A 0.8 *m/z* isolation window was typically employed for most of the targets; in special cases, 0.5-0.6 *m/z* window was used to avoid overlapping signal. MS/MS experiments were performed by scanning the nano-DESI probe over the selected region with the selected isolation window at 2.5-4 μm/s scan rate. MS/MS data acquisition was conducted in the I^2^MS mode with an Orbitrap detection period of 2 s (HCD pressure setting = 0.5) (*41*). HCD collision energy and injection time was optimized to maximize the population of individual ion fragments. Typical ranges of collision energy and injection time used in this study were 7-14 eV and 200-1500 ms, respectively. Total data acquisition time for each target varied from 1-5 hours.

MS/MS data was first subjected to I^2^MS processing for fragment ion charge assignment and mass-domain spectrum construction following the same procedure as described above. Mass-domain spectrum was converted to *.mzML* format subjected to MS^2^ search function implemented in ProSight Native (Proteinaceous, Evanston, IL) to look for possible candidates from the entire human protein database. For each search, the top 1-5 candidates were manually validated using a custom version of TDValidator (Proteinaceous, Evanston, IL) to identify the best matching proteoform. Proteoform E-values were obtained from ProSight Native and TD Validator reports.

## Funding

National Institutes of Health UH3 CA246635 (NLK)

National Institutes of Health P41 GM108569 (NLK)

National Institutes of Health P30 DA018310 (NLK)

National Institutes of Health P30 CA060553 (awarded to the Robert H. Lurie Comprehensive Cancer Center)

National Institutes of Health UH3CA255132 (JL)

National Institutes of Health U54DK120058 (JMS)

National Institute of Environmental Health Sciences T32ES007028 (EKN).

NIH National Cancer Institute 5 UM1 CA183727-08 (Cooperative Human Tissue

Network at Vanderbilt University Medical Center).

## Author contributions

Conceptualization: PS, JMC, JOK, NLK

Methodology: PS, MY, FAB, JOK, JL, NLK

Resources: MY, EKN, JLA

Software: KRD, MARH, JBG, BPE, RTF

Investigation: PS, EKN, FAB

Visualization: PS, JPM, KRD, MARH, JBG, BPE, JOK

Supervision: JMC, JOK, NLK

Writing—original draft: PS, JMC, NLK

Writing—review & editing: PS, JPM, KRB, MARH, MY, EKN, JLA, BSD, FAB,

JBG, BPE, RTF, JMC, JOK, JMS, JL, NLK

## Data and materials availability

Custom compiled code used to process and create I^2^MS files is already available *(39)*. Additional software and data that support the findings of this study are available from the corresponding authors upon request.

