## Supplementary Figures for "Direct Imaging and Identification of Proteoforms up to 70 kDa from Human Tissue"

**This PDF file includes:**

Supplementary Text

Figs. S1 to S7

Supplementary Text

Development of PiMS

Sampling conditions in PiMS is critical to imaging and identification of proteoforms. In particular, we optimized nano-DESI probe scan rate and MS injection time to achieve efficient sampling of individual ions from the tissue. In these studies, mouse brain tissue sections were selected as the model system, which has been extensively studied in imaging MS. We evaluate the sampling efficiency based off the number of protein ions obtained from a 100 µm × 150 µm unit area on the tissue. The sampling unit areas selected in this study span across all major histological regions in the brain section.

To match the data acquisition rate of PiMS experiments, nano-DESI probe scan rate was optimized in the 1-10 µm/s range. Figure 2a shows histograms of total ion count (TIC) from 100 µm × 150 µm unit sampling areas at different scan rates. TICs increase with lower scan rate (fig. S3). This is explained by the higher extraction efficiency with longer residing time and higher total volume of extraction solvent consumed for the unit sampling area. However, the standard deviation of TIC drastically increases at lowest scan rate (1 µm/s). The scans with lower TICs are a result of oversampling with a stalling liquid bridge when the probe moves at extremely low scan rate. The oversampling phenomena are evidenced by consecutive regions of exponentially decayed total ion chronogram of the line acquisition (fig. S4). Considering the sampling continuity, efficiency, and experimental throughput, we choose 2.5-5 µm/s as the typical scan rate used for subsequent PiMS experiments.

MS injection time is another contributing factor to the individual ion population obtainable from a unit sampling area on the tissue. We first performed I^2^MS processing on the line scans obtained at different MS injection times, which converted the m/z of the protein ions into the absolute mass domain. Fig. S5 shows the mass-domain proteoform landscape of a mouse brain tissue section. In addition to low mass proteoforms <20 kDa, we observe a broad landscape of proteoforms up to 80 kDa with many of them exceeding 40 kDa. Different MS injection times was found to have a pronounced effect on the mass landscape in a singular unit area (100 µm × 150 µm). Fig. S6a and S6b show the ion count statistics of different mass ranges from sampling unit areas from distinct histological regions (neocortex and cerebellum). <20 kDa proteoforms dominate the mass spectrum when injection time is relatively low. In contrast, longer injection times favor the detection of higher mass proteins with better constructed statistics. For the subsequent experiments focusing on high mass tissue proteoforms, a typical MS injection time of 200-300 ms were employed to maximize the sensitivity of the higher mass range (>20 kDa).

**Figure S1.**

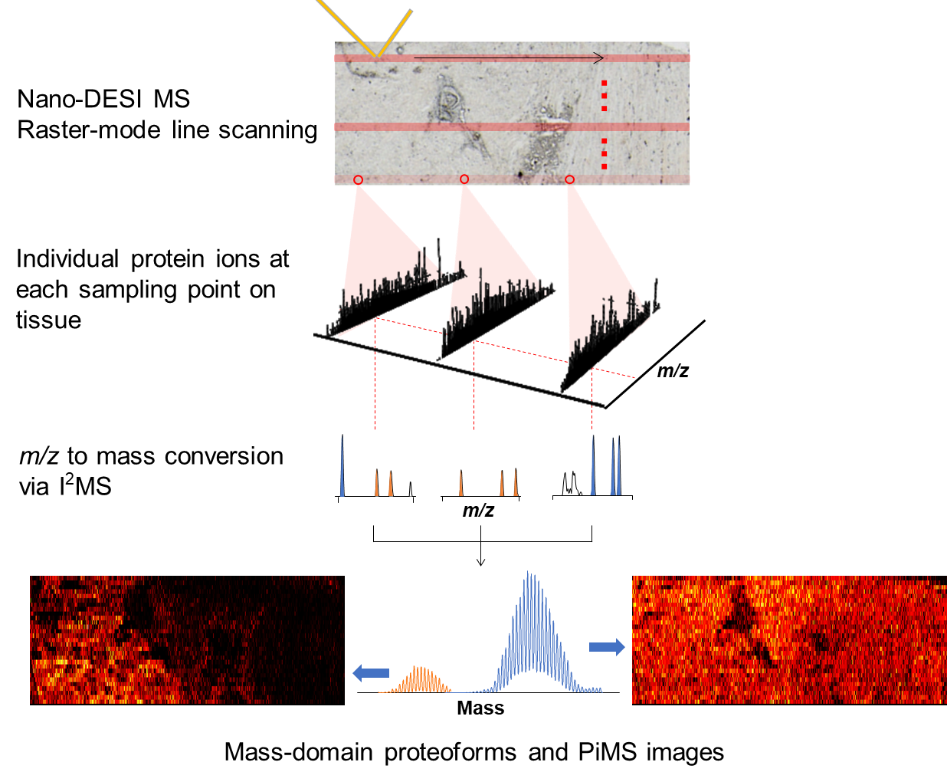

**Figure S1.** The PiMS workflow. The schematic shows the scanning approach (top), collection and interpretation of individual ions into the mass domain (middle) and image reconstruction (bottom). Representative MS full spectra of singular scans are displayed in the data collection step, demonstrating the collection of individual ions from different locations on the tissue. The I^2^MS processing pipeline has been previously published and described in detail in the Method section.

**Figure S2.**

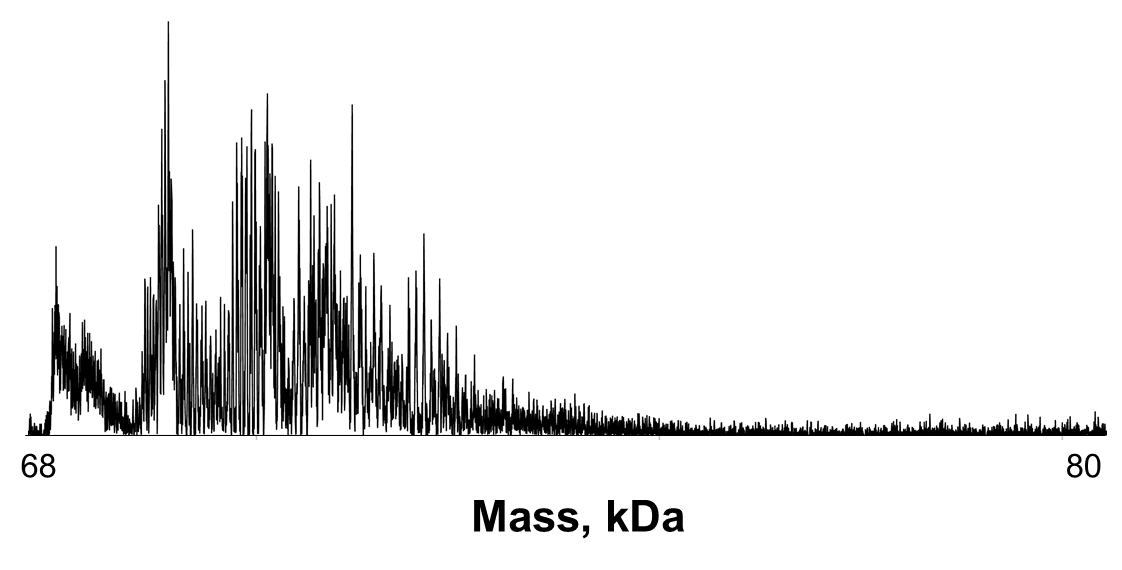

**Figure S2.** Full PiMS spectrum in the mass range of 68-80 kDa showing protein signals not individually resolved.

**Figure S3.**

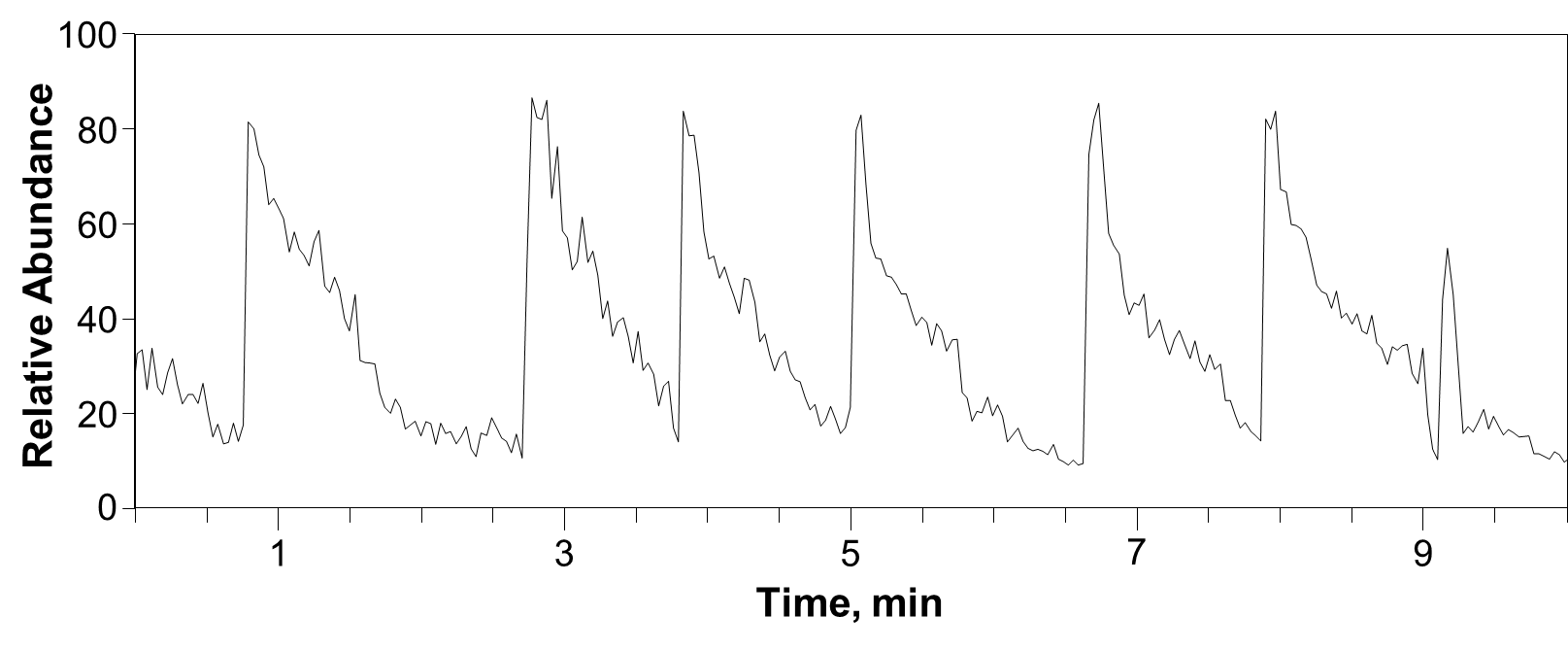

**Figure S3.** Total ion count as a function of time during part of a PiMS line scan at 1 µm/s with exponentially decaying profiles, indicating a stalling liquid bridge when the probe moves at extremely low scan rate.

**Figure S4.**

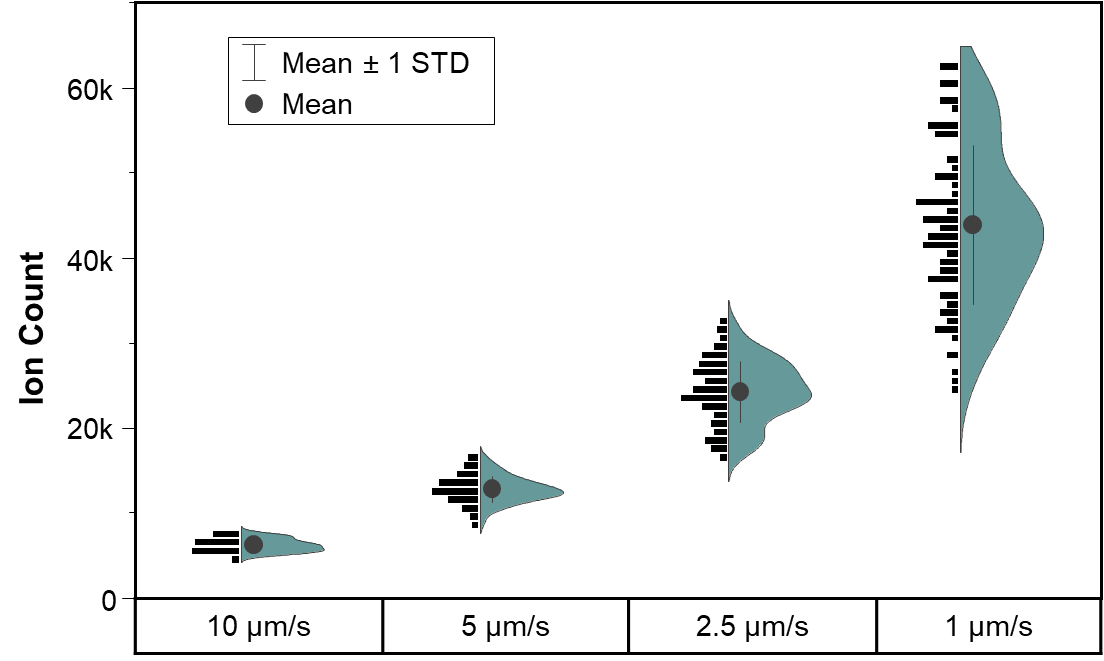

**Figure S4.** Individual ion statistics from 100 µm × 150 µm areas on a mouse brain tissue section at different scan rate at 50 ms injection time during PiMS. The plots consist only of ions passing the STORI analyses used in I^2^MS and recognized as individual ions. The violin plots mirrored to the right are Kernel Density Estimated versions of the histograms.

**Figure S5.**

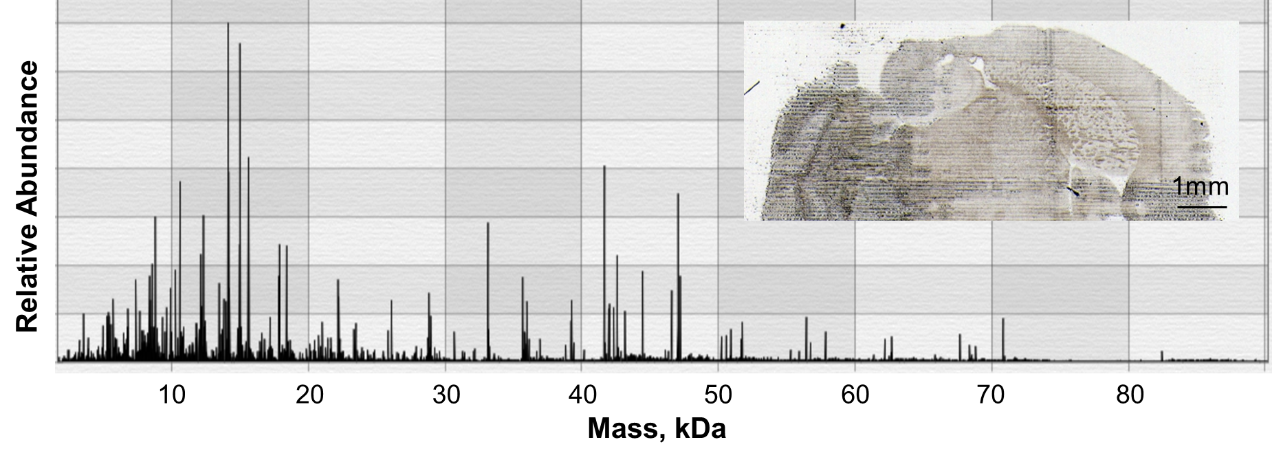

**Figure S5.** PiMS spectrum obtained from integration of individual spectra from a single line scan run at 4 µm/s across a mouse brain tissue section; (inset, optical image of the tissue section after PiMS experiment).

**Figure S6.**

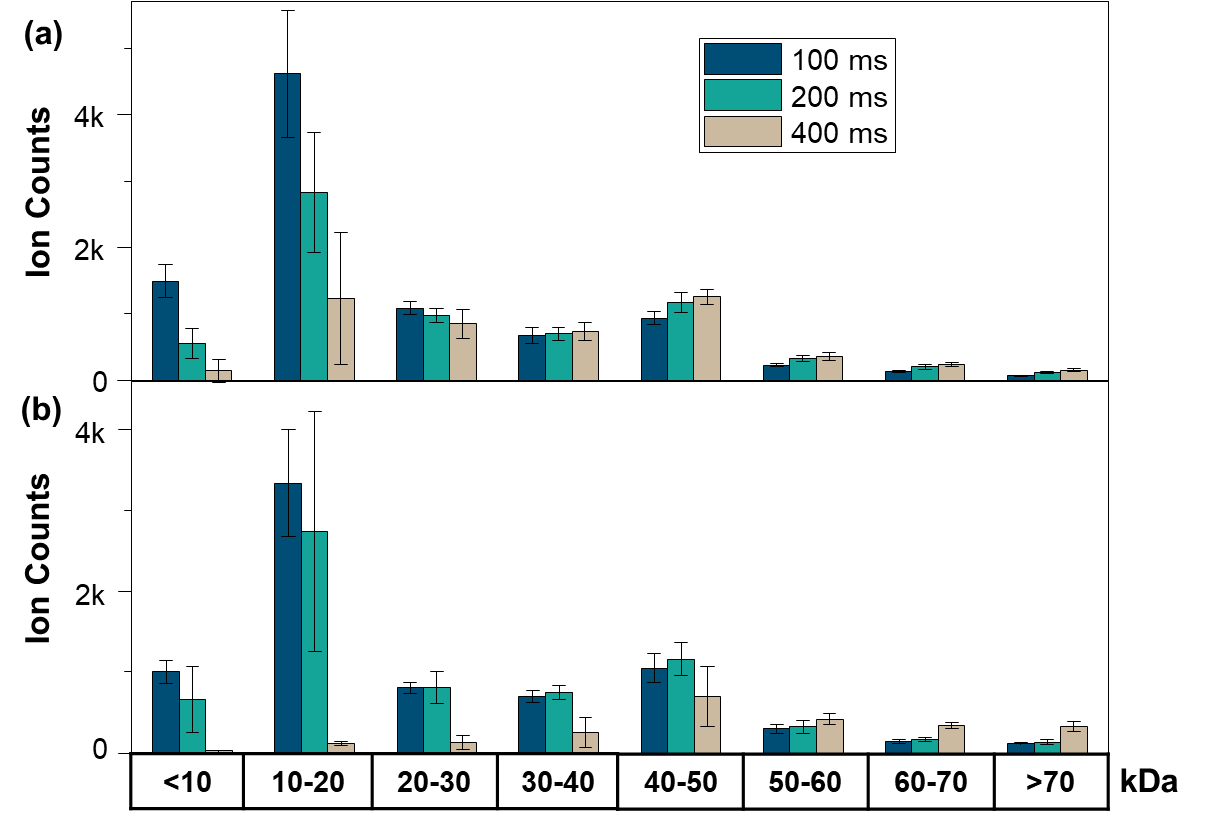

**Figure S6.** Raw ion counts obtained at three ion injection times during PiMS of mouse brain tissue running at a scan rate of 4 µm/s. The bar graphs show the averaged total ion counts from 100 unit areas of 100 µm × 150 µm size in the neocortex (a) and cerebellum (b) regions. Error bars indicate the standard deviation of 100 trials.

**Figure S7. Tandem mass spectra and graphical fragment maps used in the identification of 21 proteoforms via on-tissue top-down MS/MS.** Panels **a-u** show the data for the individual MS/MS experiments used for proteoform identification.

**(a) Alpha crystallin**

Precursor mass: 20201.2 Da

Isolation window (charge state): 595.1 ±0.25 (34+)

Accession: P02511

Sequence coverage: 35%

E-value: 1.1E-131

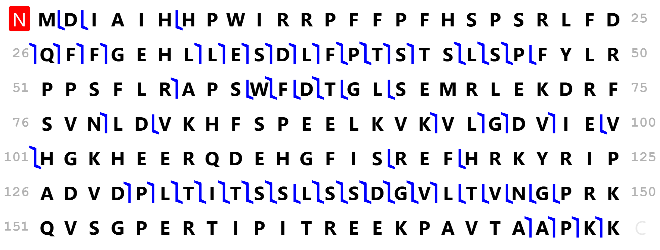

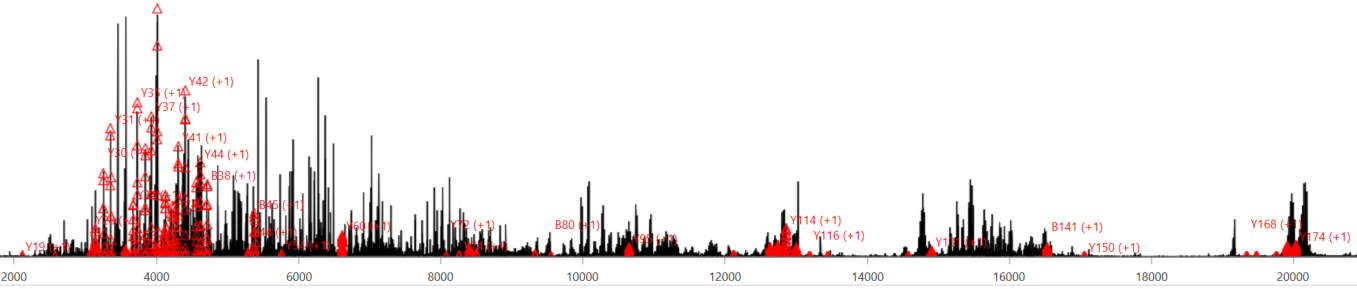
 **Mass, Da**

**(b) Superoxide dismutase [Mn], mitochondrial**

Precursor mass: 22204.3 Da

Isolation window (charge state): 926.1±0.3 (24+)

Accession: P04179-1

Sequence coverage: 16%

E-value: 2.1E-58

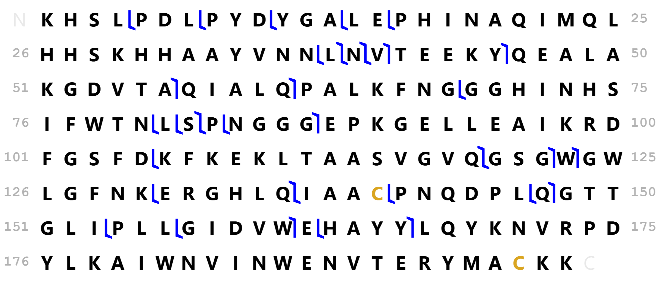

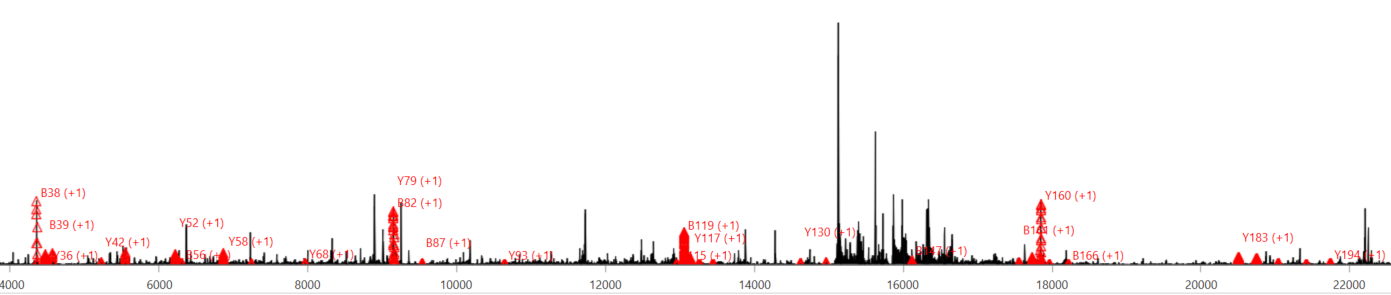
**Mass, Da**

**(c) Membrane-spanning 4-domains subfamily A member 4A**

Precursor mass: 25427.5 Da

Isolation window (charge state): 979±0.4 (26+)

Accession: Q96JQ5

Sequence coverage: 4%

E-value: 8.3E-18

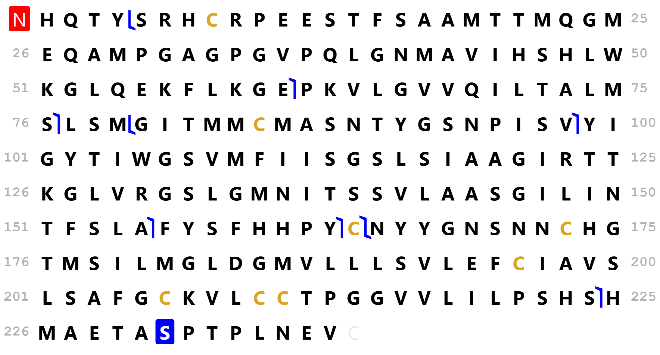

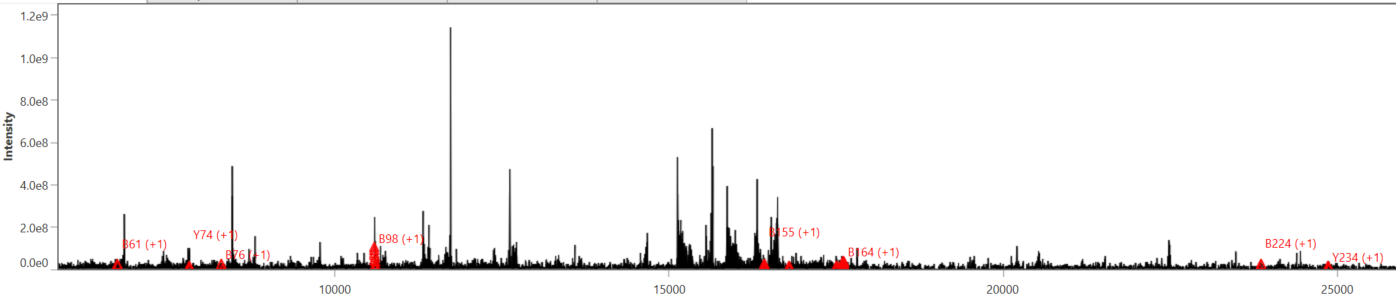
**Mass, Da**

**(d)**

**Glutathione S-transferase A2 (canonical sequence)**

Precursor mass: 25588.6 Da

Isolation window (charge state): 914.9±0.2 (28+)

Accession: P09210

Sequence coverage: 24%

E-value: 8.6E-87

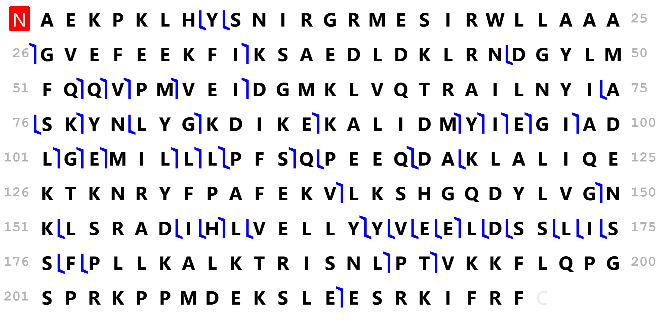

**Glutathione S-transferase A2 (Ser111**→**Thr)**

Precursor mass: 25574.6 Da

Isolation window (charge state): 914.9±0.2 (28+)

Accession: P09210

Sequence coverage: 24%

E-value: 3.1E-87

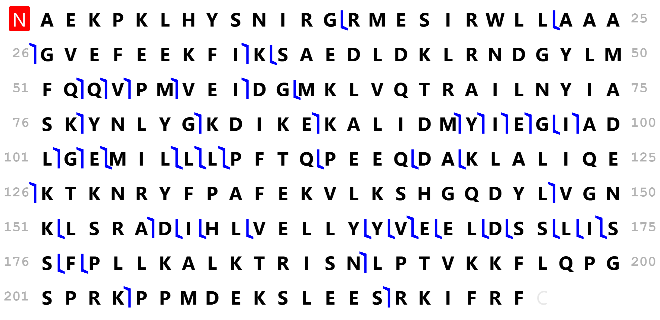

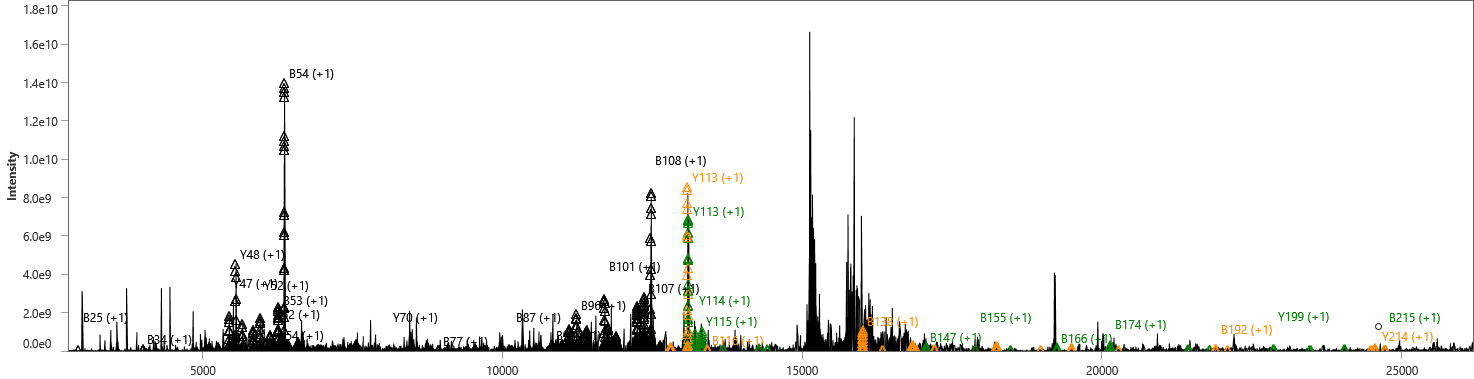
**Mass, Da**

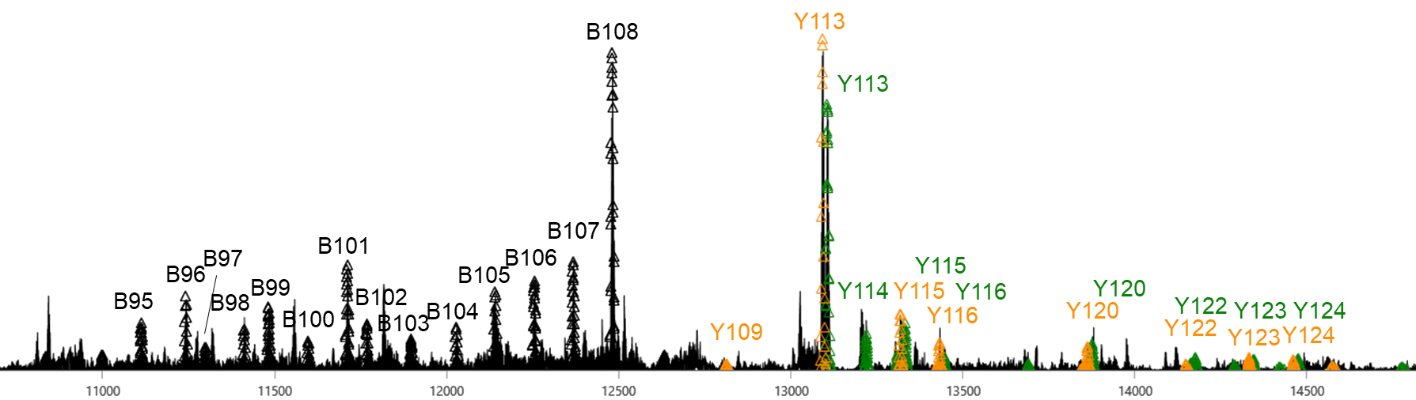
 **Mass, Da**

Fragments in yellow indicate canonical sequence, fragments in green indicate Ser111→Thr natural variant, and fragments in black indicate common fragments.

**(e)**

**V-type proton ATPase subunit E 1**

Precursor mass: 26055.7 Da

Isolation window (charge state): 815.2±0.2 (32+)

Accession: P36543-1

Sequence coverage: 14%

E-value: 4.2E-54

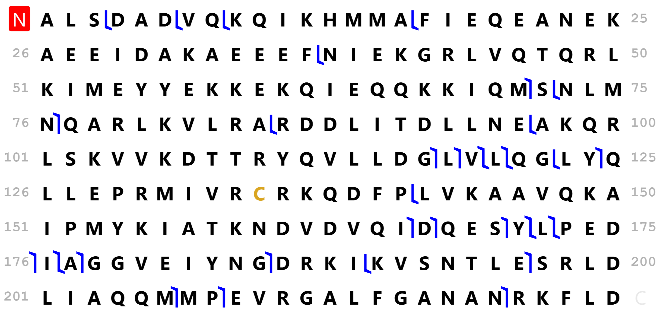

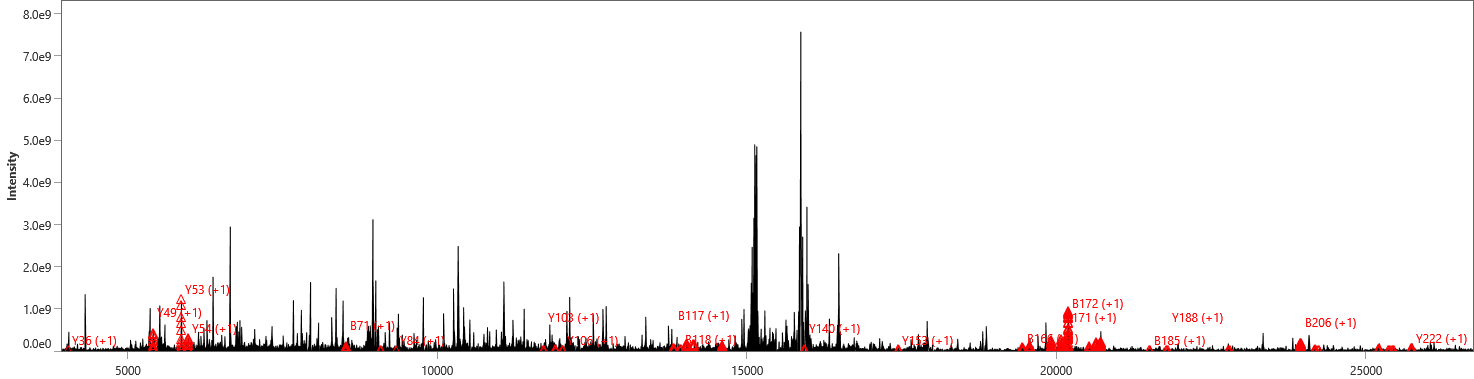
 **Mass, Da**

**(f)**

**Apolipoprotein A-1**

Precursor mass: 28078.4 Da

Isolation window (charge state): 780.9±0.3 (36+)

Accession: P02647

Sequence coverage: 17%

E-value: 1.7E-65

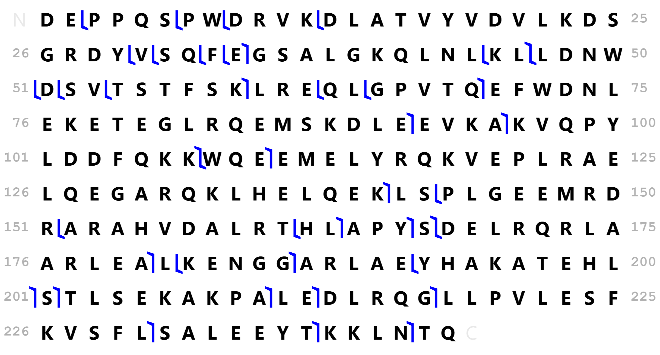

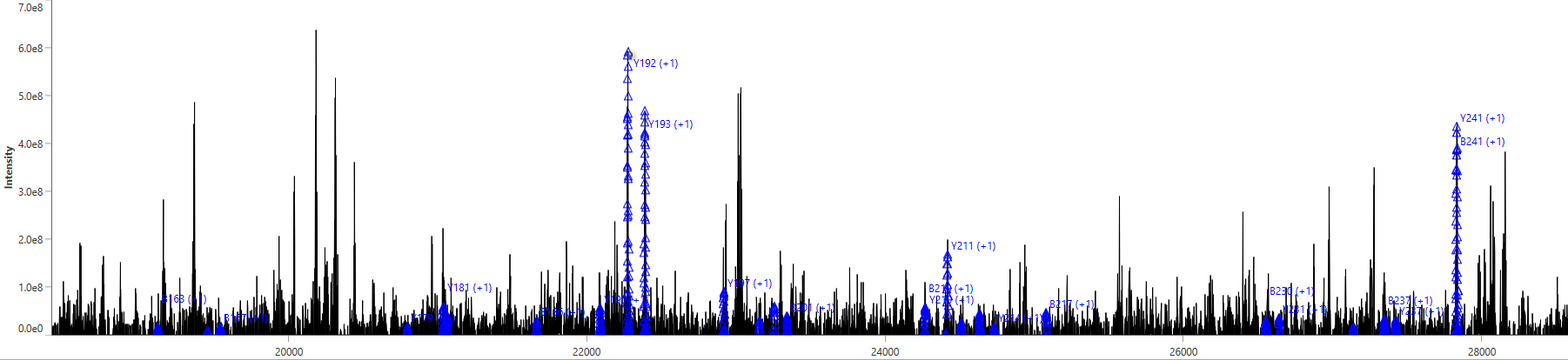

**Mass, Da**

**(g)**

**Adaptin ear-binding coat-associated protein 1**

Precursor mass: 29766.0 Da

Isolation window (charge state): 784.4±0.4 (38+)

Accession: Q8NC96-1

Sequence coverage: 18%

E-value: 3.5E-75

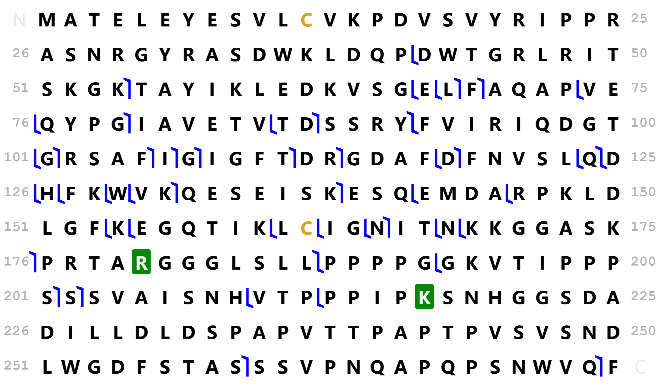

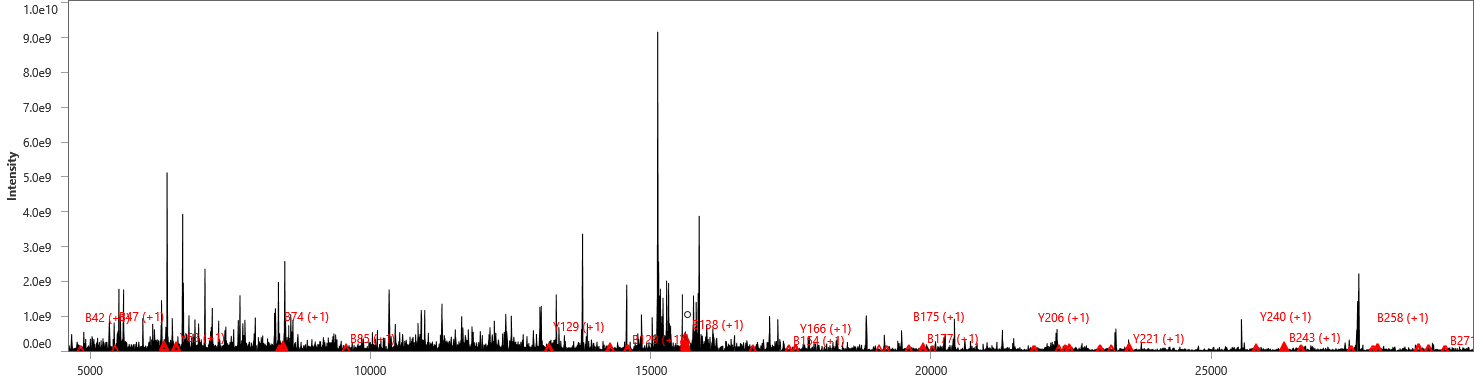
 **Mass, Da**

**(h)**

**N(G),N(G)-dimethylarginine dimethylaminohydrolase 1**

Precursor mass: 31032.9 Da

Isolation window (charge state): 863.0±0.3 (36+)

Accession: O94760

Sequence coverage: 16%

E-value: 1.3E-74

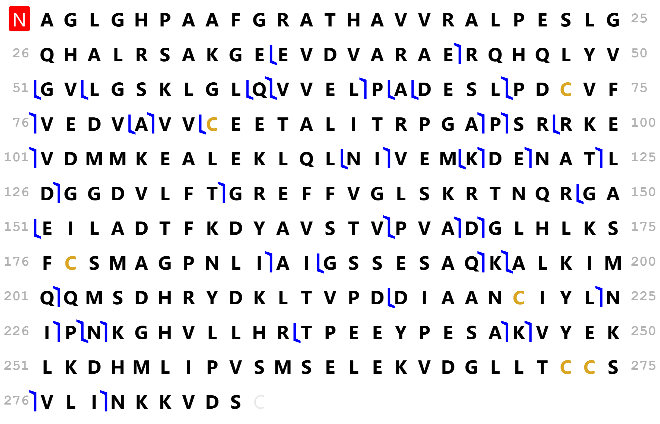

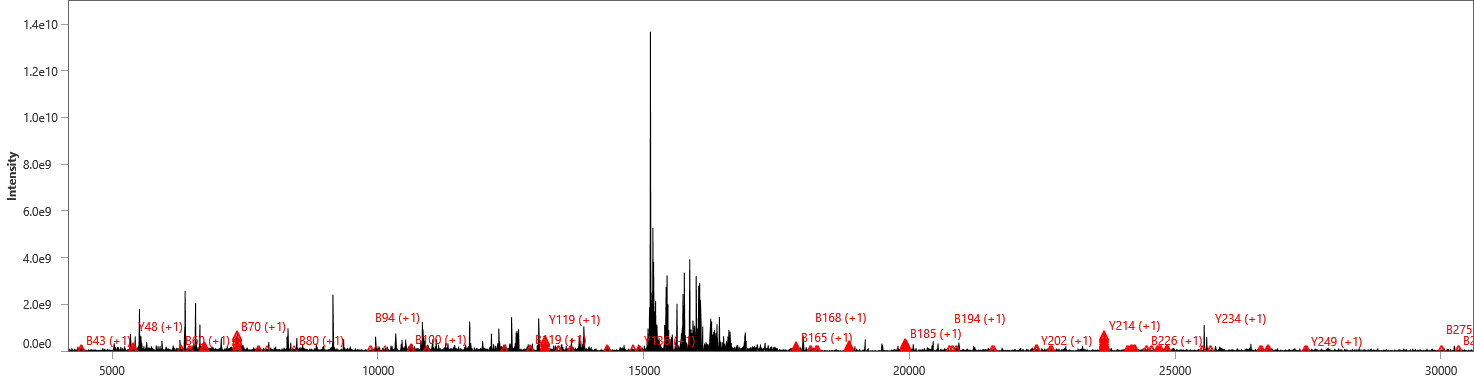
 **Mass, Da**

**(j)**

**Malate dehydrogenase, mitochondrial**

Precursor mass: 33000.4 Da

Isolation window (charge state): 1001.0±0.2 (33+)

Accession: P40926-1

Sequence coverage: 18%

E-value: 6.1E-94

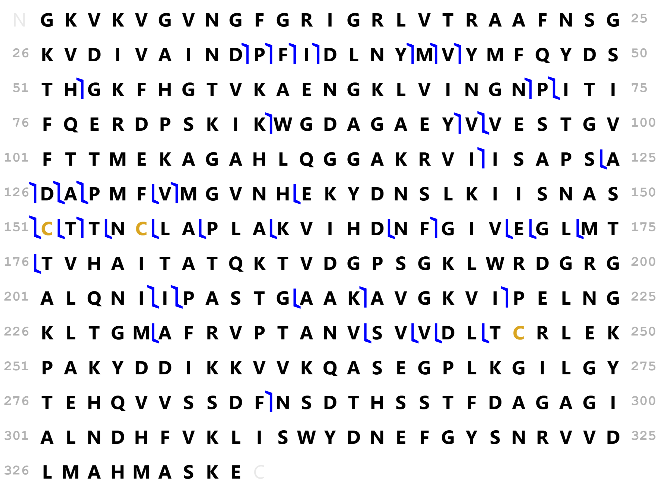

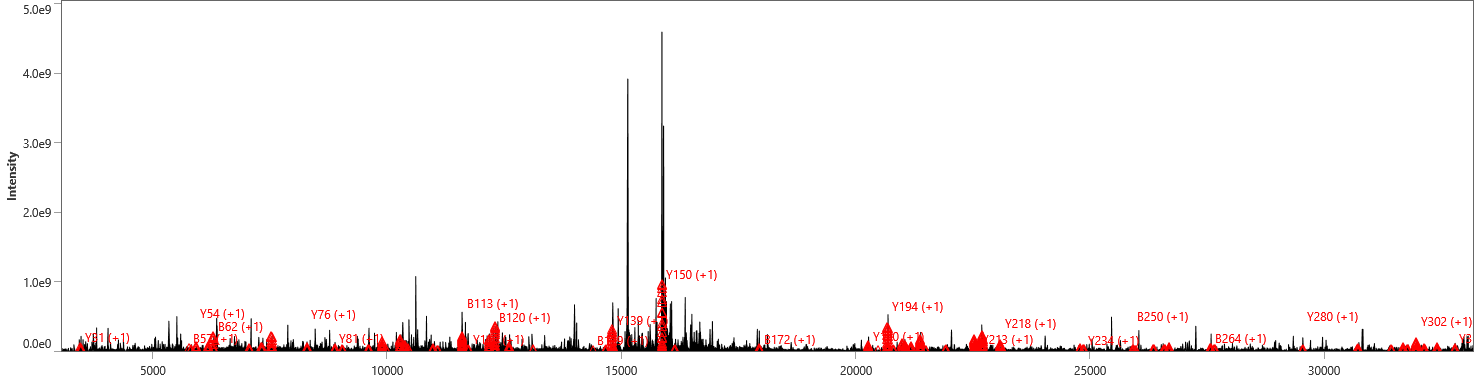
 **Mass, Da**

**(k)**

**Glyceraldehyde-3-phosphate dehydrogenase**

Precursor mass: 35922.4 Da

Isolation window (charge state): 781.9±0.4 (46+)

Accession: P04406-1

Sequence coverage: 14%

E-value: 4.5E-76

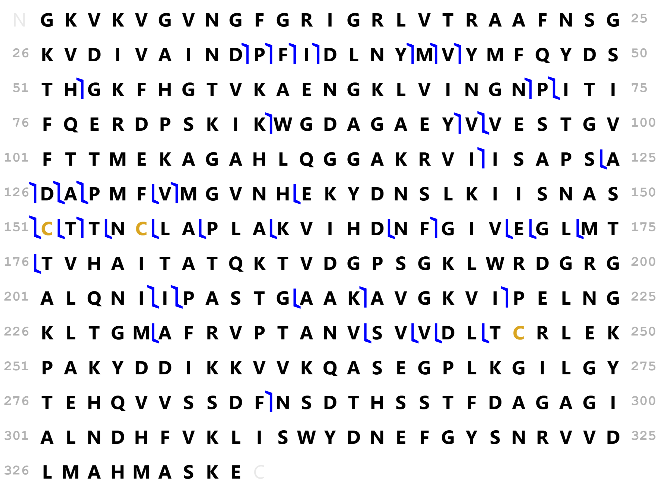

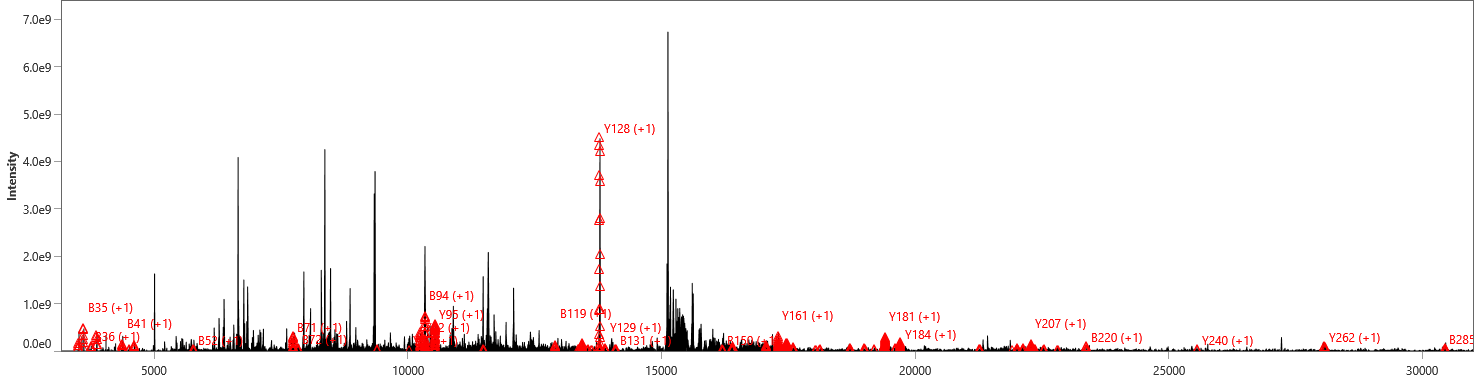
 **Mass, Da**

**(l)**

**Glyceraldehyde-3-phosphate dehydrogenase**

Precursor mass: 35951.4 Da

Isolation window (charge state): 922.9±0.2 (39+)

Accession: P04406-1

Sequence coverage: 14%

E-value: 8.6E-63

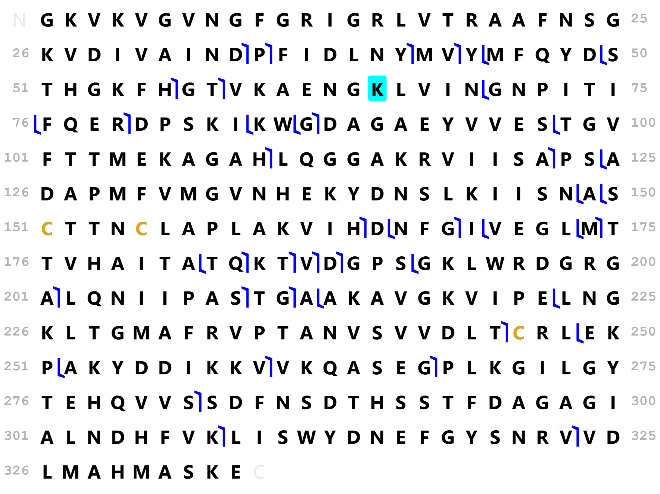

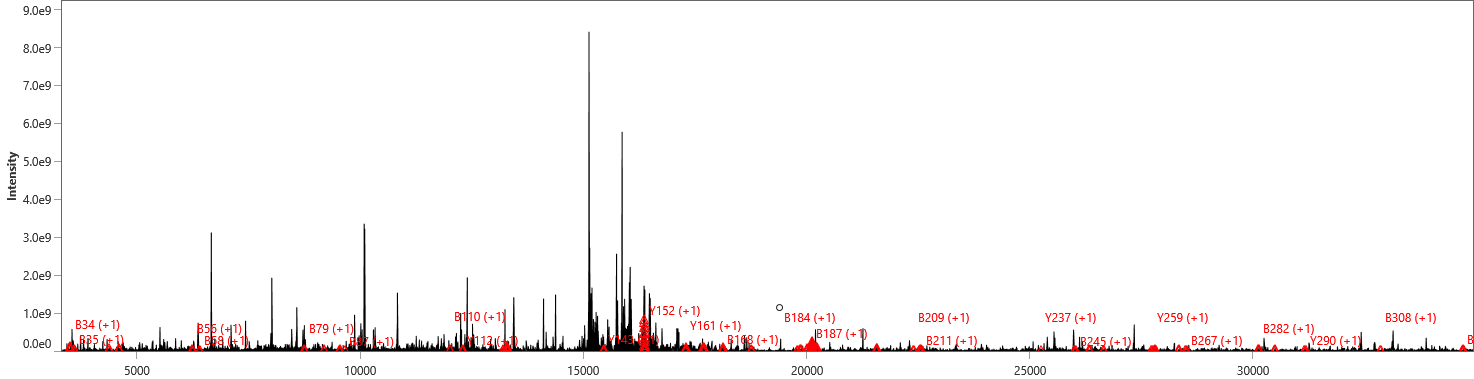
 **Mass, Da**

**(m)**

**Fructose-1,6-bisphosphatase 1**

Precursor mass: 36725.7 Da

Isolation window (charge state): 967.4±0.2 (38+)

Accession: P09467

Sequence coverage: 22%

E-value: 1.1E-121

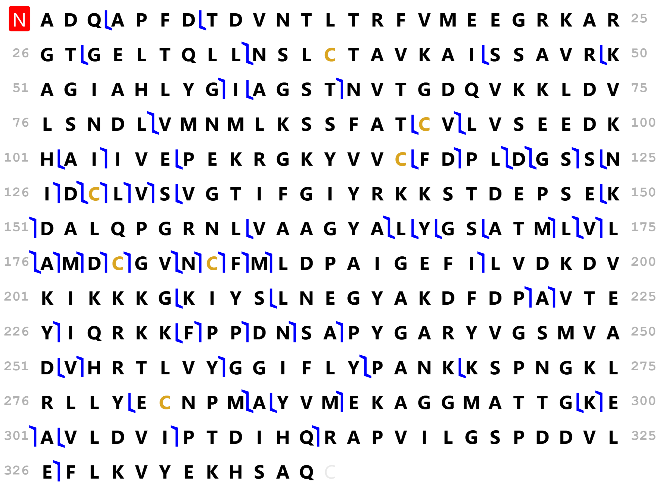

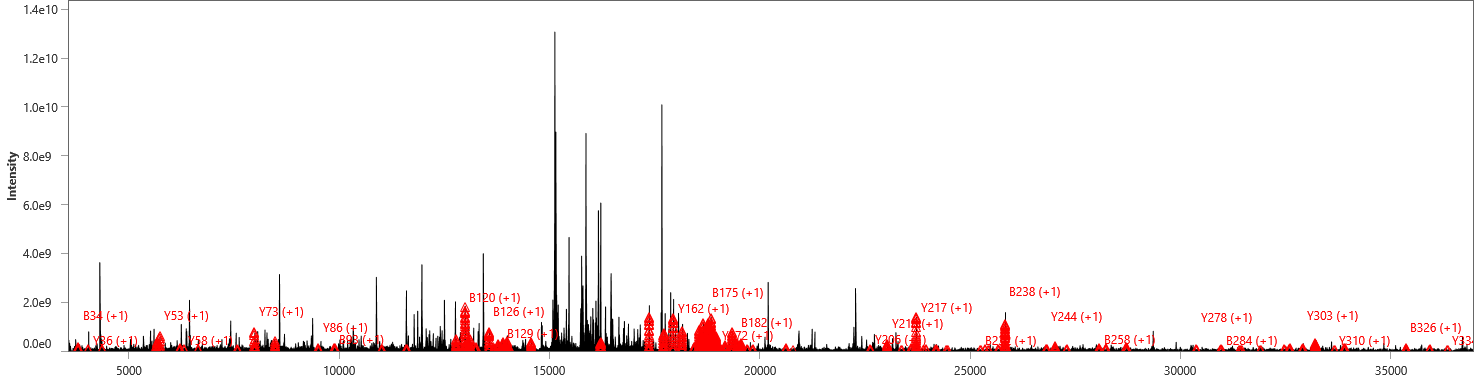
 **Mass, Da**

**(n)**

**Actin, cytoplasmic 1**

Precursor mass: 41661.7 Da

Isolation window (charge state): 969.9 ±0.2 (43+)

Accession: P60709

Sequence coverage: 24%

E-value: 3.8E-160

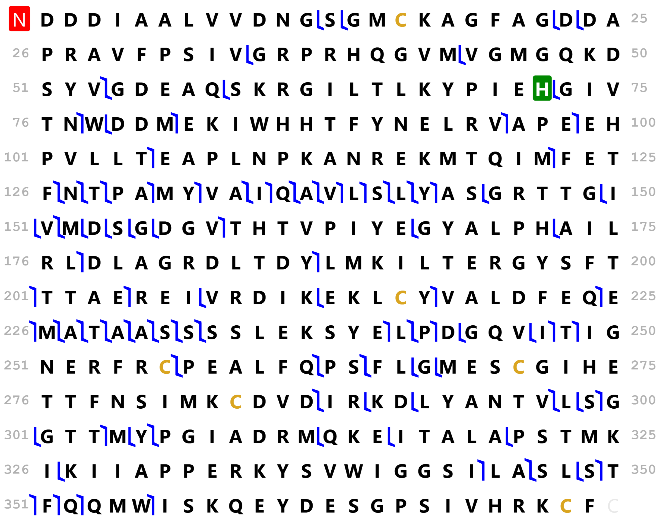

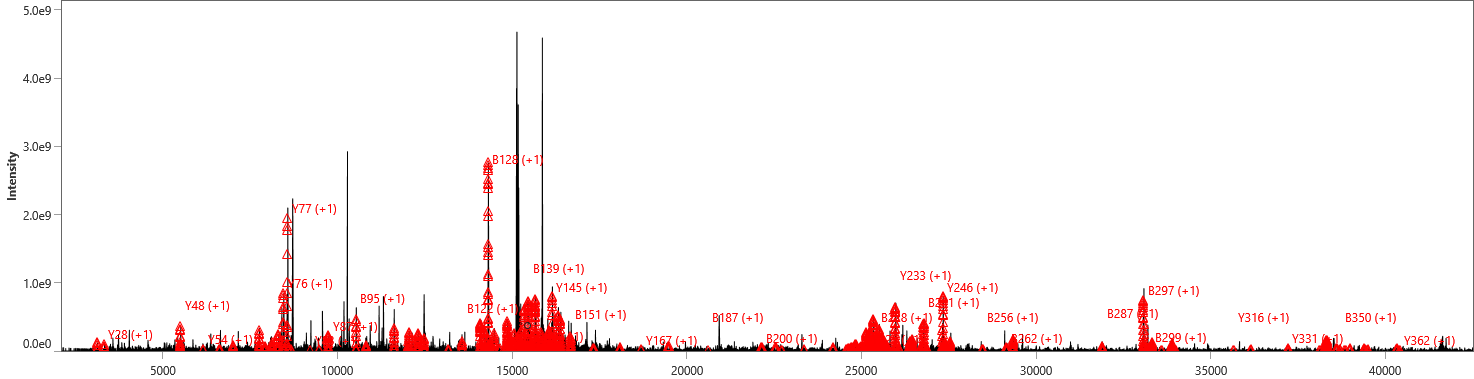
**Mass, Da**

**(p)**

**Actin, cytoplasmic 2**

Precursor mass: 41661.7 Da

Isolation window (charge state): 969.9±0.2 (43+)

Accession: P63261

Sequence coverage: 24%

E-value: 1.1E-154

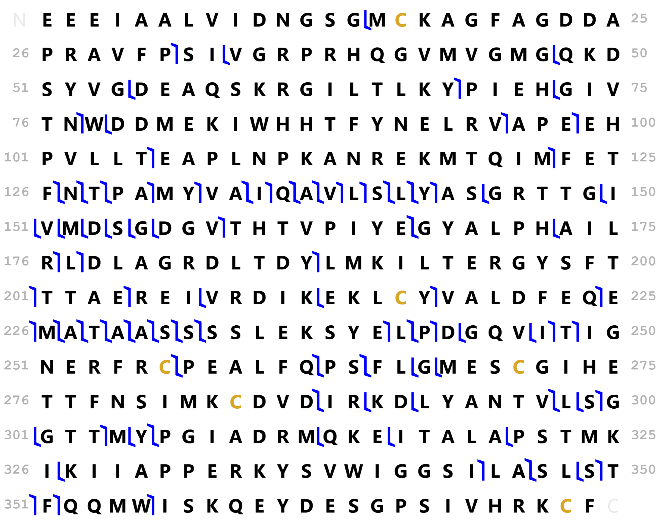

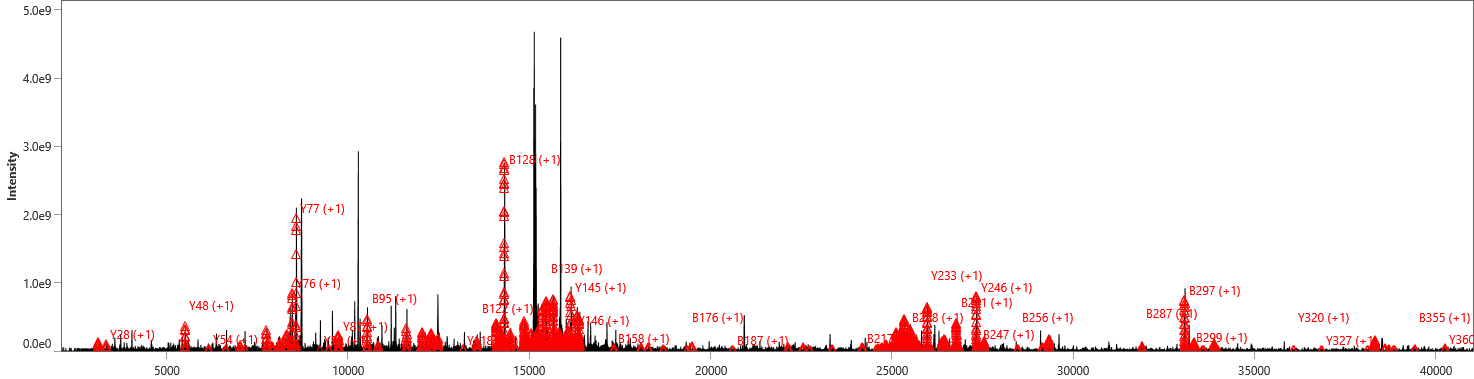
**Mass, Da**

**(q)**

**Actin, cytoplasmic 2**

Precursor mass: 41716.8 Da

Isolation window (charge state): 1043.9±0.2 (40+)

Accession: P63261

Sequence coverage: 20%

E-value: 1.7E-115

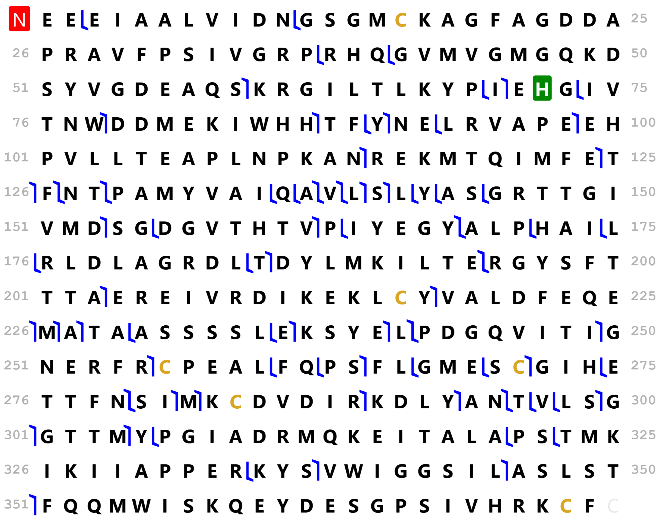

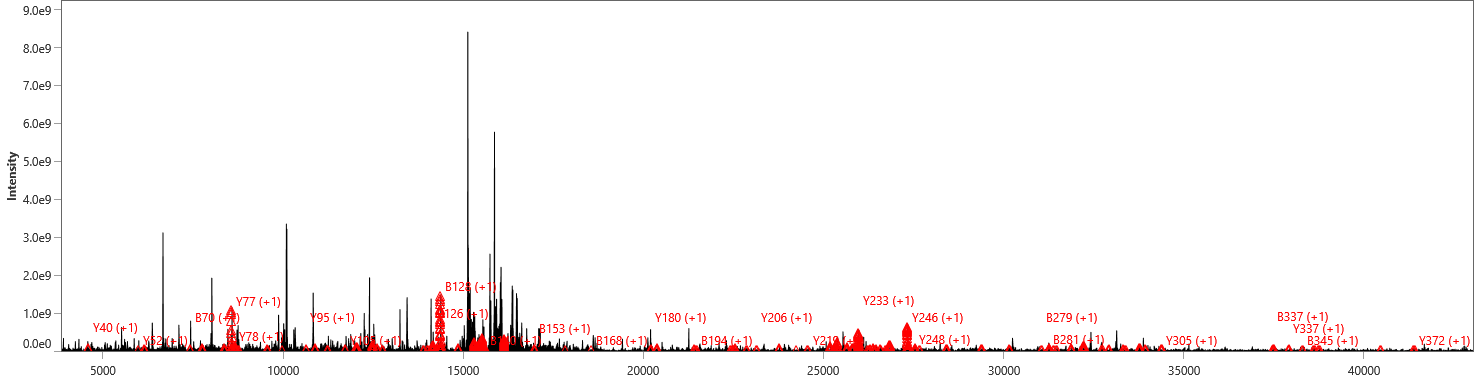
 **Mass, Da**

**(r)**

**Alpha-enolase**

Precursor mass: 47078.8 Da

Isolation window (charge state): 981.8±0.4 (48+)

Accession: P06733

Sequence coverage: 12%

E-value: 1.6E-74

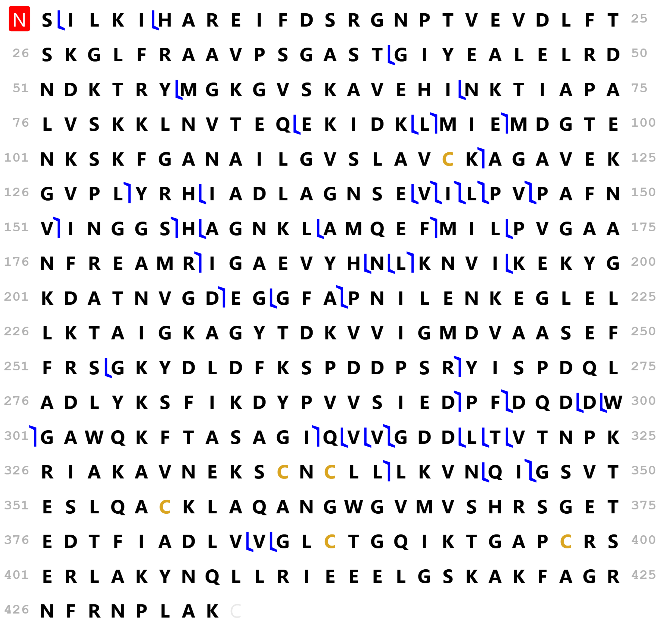

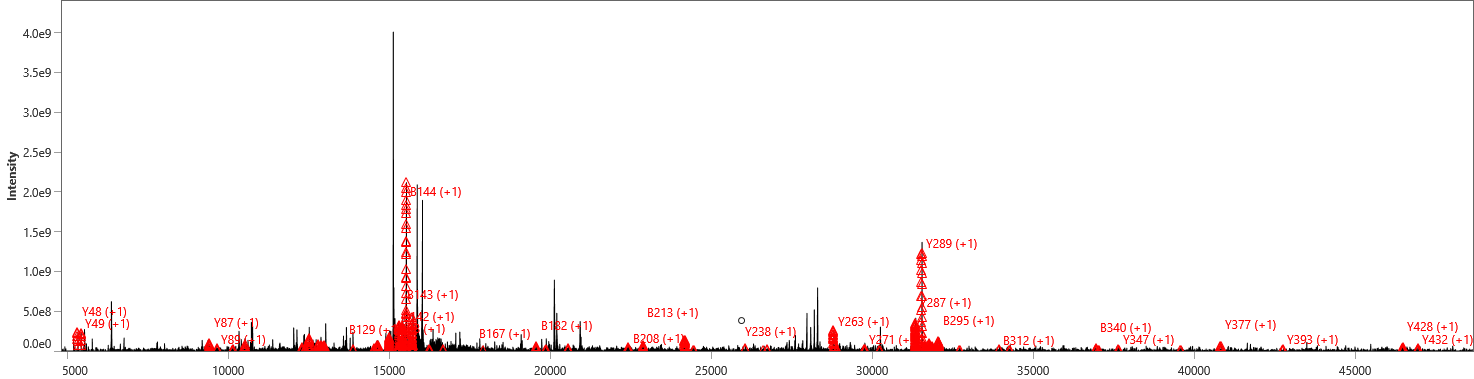
**Mass, Da**

**(s)**

**Vimentin**
Precursor mass: 53561.5 Da

Isolation window (charge state): 851.2±0.2 (63+)

Accession: P08670

Sequence coverage: 16%

E-value: 4.8E-106

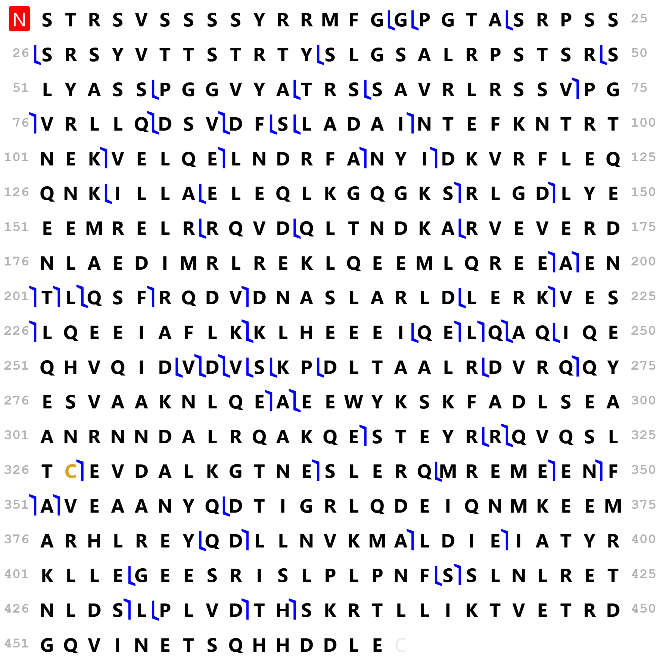

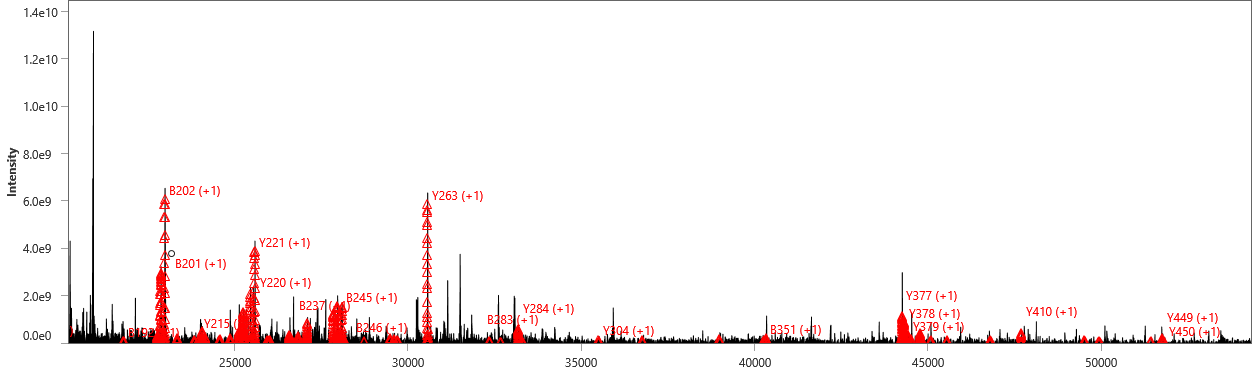
 **Mass, Da**

**(t)**

**Albumin**
Precursor mass: 66435.9 Da

Isolation window (charge state): 1208.9±0.4 (55+)

Accession: P02768

Sequence coverage: 3%

E-value: 8.3E-22

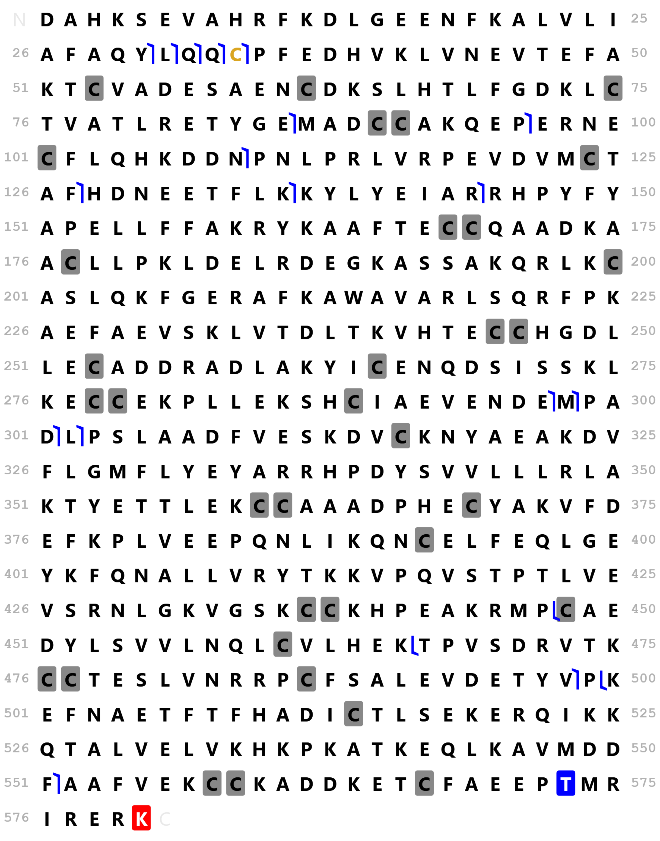

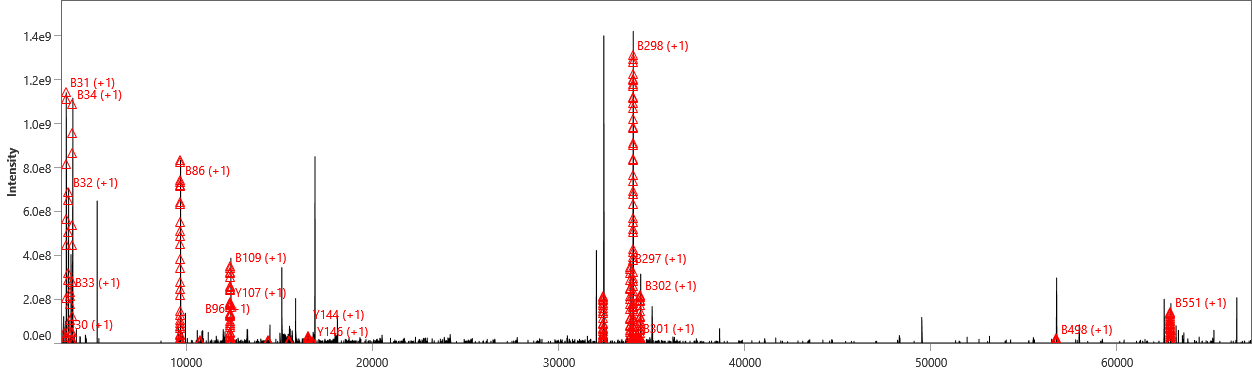
**Mass, Da**

**(u)**

**Mesothelin**
Precursor mass: 70900.52 Da

Isolation window (charge state): 1209.6±0.3 (58+)

Accession: Q13421-3

Sequence coverage: 7%

E-value: 1.60E-45

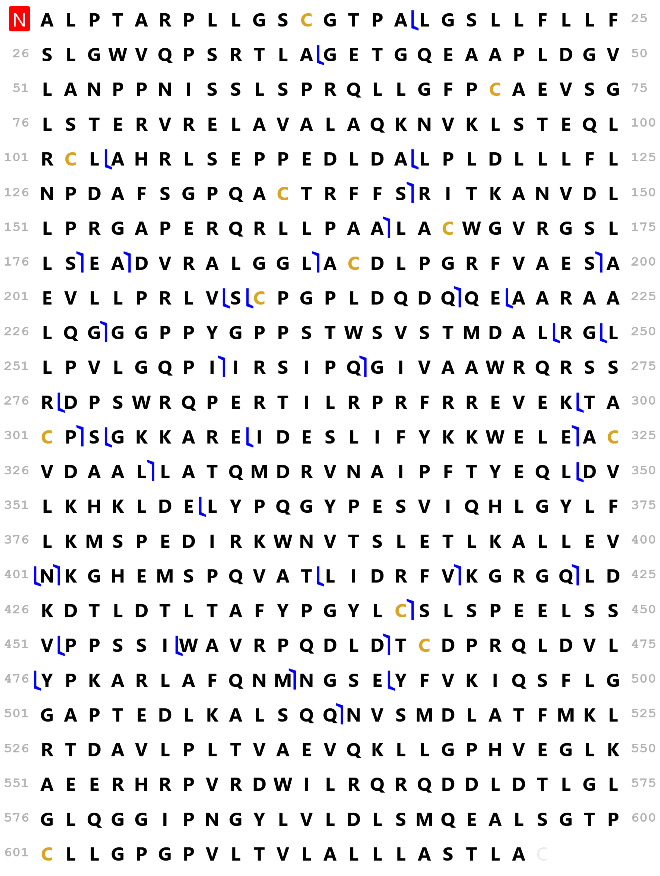

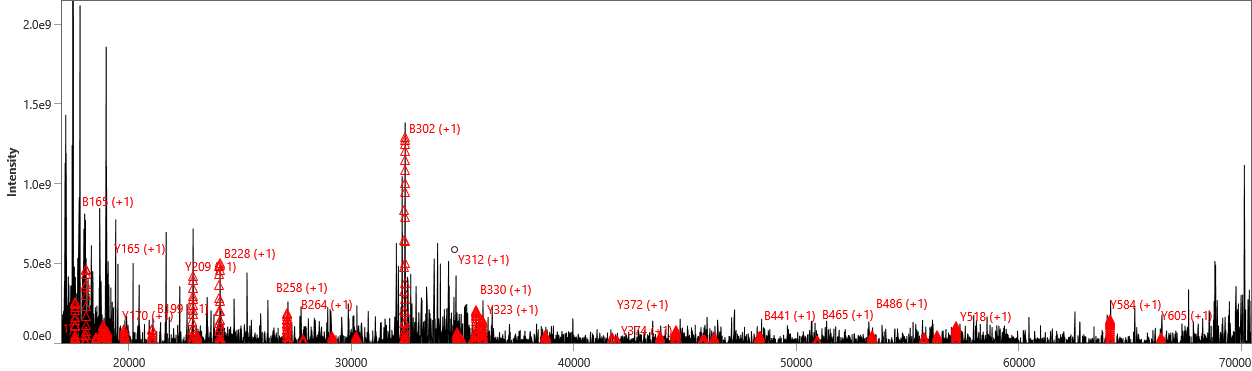
 **Mass, Da**

**Figure S7.** Technical replicates of PiMS images of adjacent kidney tissue sections from the same subject.

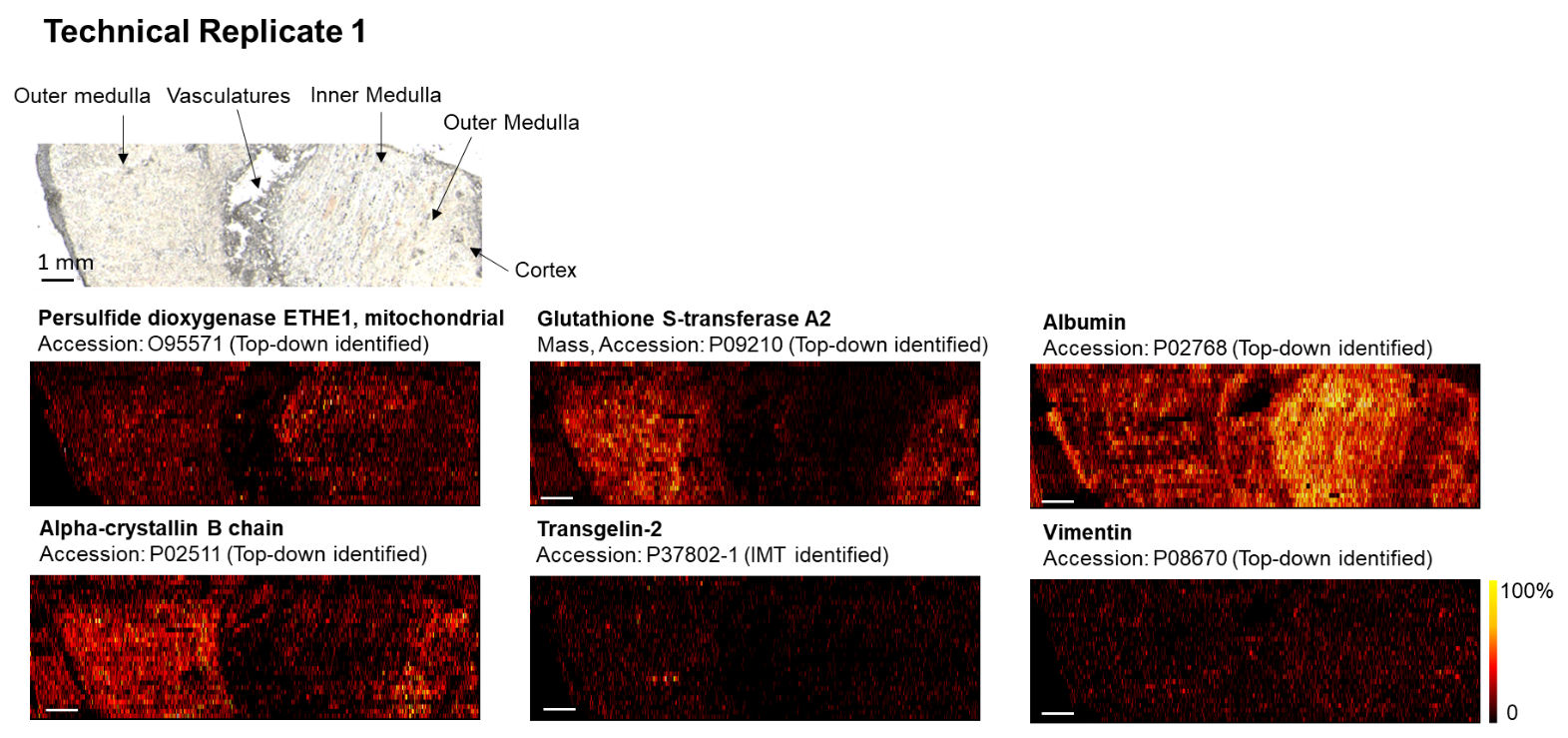

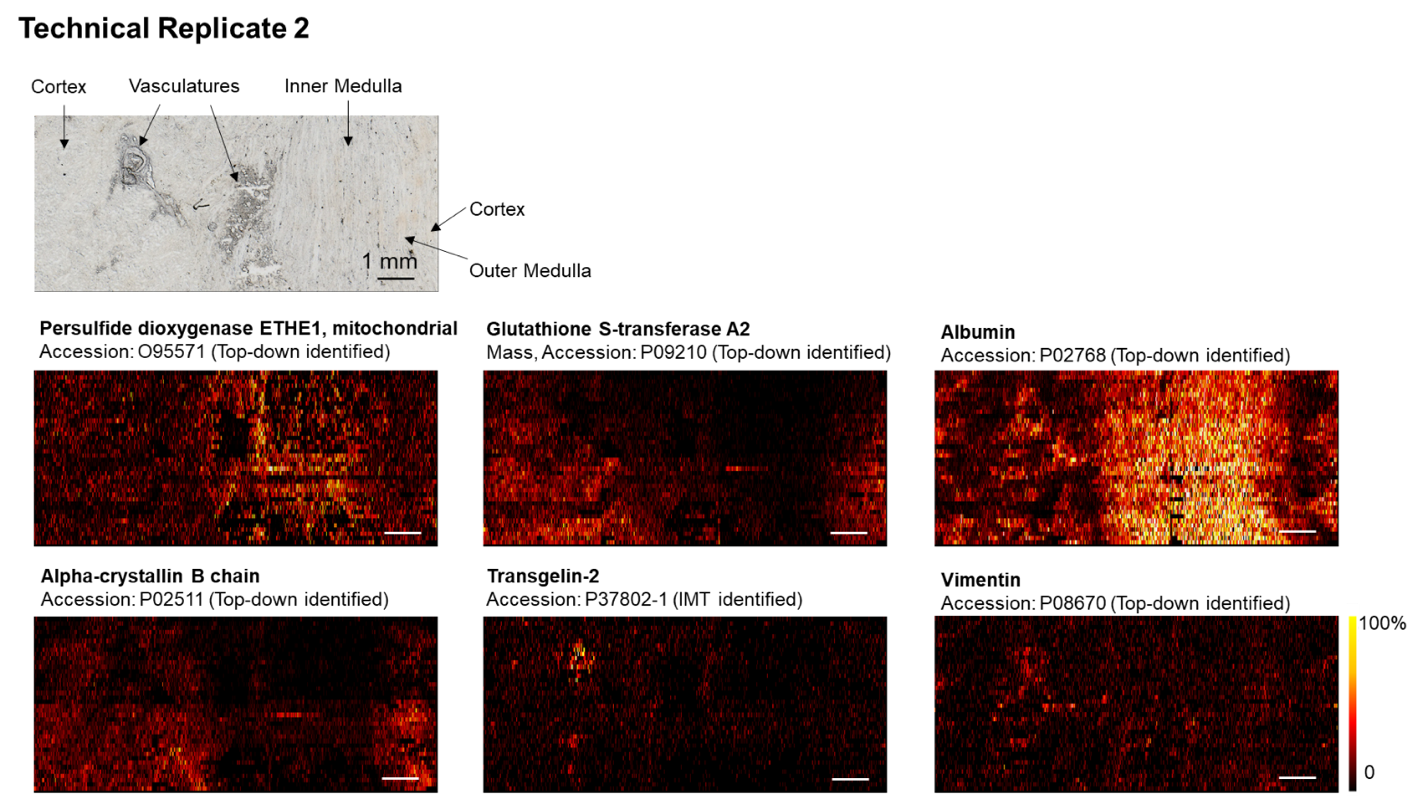
